# Functional genomics of lipid metabolism in the oleaginous yeast *Rhodosporidium toruloides*

**DOI:** 10.1101/190546

**Authors:** Samuel T Coradetti, Dominic Pinel, Gina Geiselman, Masakazu Ito, Stephen Mondo, Morgann C Reilly, Ya-Fang Cheng, Stefan Bauer, Igor V Grigoriev, John M Gladden, Blake A Simmons, Rachel B Brem, Adam P Arkin, Jeffrey M Skerker

## Abstract

The basidomycete yeast *Rhodosporidium toruloides* (a.k.a. *Rhodotorula toruloides)* accumulates high concentrations of lipids and carotenoids from diverse carbon sources. It has great potential as a model for the cellular biology of lipid droplets and for sustainable chemical production. We developed a method for high-throughput genetics (RB-TDNAseq), using sequence-barcoded *Agrobacterium tumefaciens* T-DNA insertions into the *R. toruloides* genome. We identified 1337 putative essential genes with low T-DNA insertion rates. We functionally profiled genes required for fatty acid catabolism and lipid accumulation, validating results with 35 targeted deletion strains. We found that both mitochondrial and peroxisomal enzymes were required for growth on fatty acids, with different peroxisomal enzymes required on different fatty acids. We identified a high-confidence set of 150 genes affecting lipid accumulation, including genes with predicted function in signaling cascades, gene expression, protein modification and vesicular trafficking, autophagy, amino acid synthesis and tRNA modification, as well as genes of unknown function. These results greatly advance our understanding of lipid metabolism in this oleaginous species, identify key biological processes to be further explored and optimized for production of lipid-based bioproducts, and demonstrate a general approach for barcoded mutagenesis that should enable functional genomics in diverse fungi.

## Introduction

*Rhodosporidium toruloides* (also known as *Rhodotorula toruloides* (1)) is a basidomycete yeast (subdivision Pucciniomycotina). *Rhodotorula/Rhodosporidium* species are widely distributed in the phyllosphere and diverse soils (2-5). They accumulate high concentrations of carotenoid pigments (6, 7), giving their colonies a distinctive orange, red, or pink hue. When *R. toruloides* is cultured under nitrogen (8), sulfur (9), or phosphorus (10) limitation, it can accumulate as much as 70% of cellular biomass as lipids (11), primarily as triacylglycerides (TAG).

Eukaryotes accumulate neutral lipids in complex, dynamic organelles called lipid droplets. Lipid droplets emerge from the endoplasmic reticulum (ER) membrane as a core of TAG surrounded by sterol esters, a phospholipid monolayer derived from ER phospholipids, and a targeted ensemble of proteins mediating inter-organelle interaction, protein trafficking, cellular lipid trafficking and regulated carbon flux in and out of the lipid droplet (12-14). Aberrant lipid droplet formation contributes to many human diseases (15, 16) and impacts cellular processes as diverse as autophagy (17) and mitosis (18). *R. toruloides’* propensity to form large lipid droplets under a variety of conditions makes it an attractive platform to study conserved aspects of the cellular biology of these important organelles across diverse eukaryotes.

*R. toruloides* is also an attractive host for production of sustainable chemicals and fuels from low-cost lignocellulosic feedstocks. Wild isolates of *R. toruloides* can produce lipids and carotenoids from a wide variety of carbon sources including glucose (19, 20), xylose (11), and acetate (21), as well as complex biomass hydrolysates (22). They are also relatively tolerant to many forms of stress including osmotic stress (23) and common inhibitors found in hydrolysates produced by common biomass deconstruction technologies, such as dilute acid pretreatment followed by enzymatic saccharification (24, 25). *R. toruloides* has been engineered to produce modified products such as fatty alcohols (26) and eurcic acid (27) from synthetic pathways, demonstrating this species potential for production of diverse bioproducts. To enable more efficient production of terpene-derived and lipid-derived chemicals in general, *R. toruloides* has also been engineered for enhanced carotenoid (28) and lipid (8) production. These efforts, while promising, have for the most part employed strategies adapted from those demonstrated in evolutionarily distant species such as *Saccharomyces cerevisiae* and *Yarrowia lipolytica.* To truly tap the biosynthetic potential of *R. toruloides*, a better understanding of the unique aspects of its biosynthetic pathways, gene regulation and cellular biology will be required.

Recently, transcriptomic and proteomic analysis of *R. toruloides* in nitrogen limited conditions (29) identified over 2,000 genes with altered transcript abundance and over 500 genes with altered protein abundance during lipid accumulation. These genes included many enzymes involved in the TCA cycle, a putative PYC1/MDH2/Malic Enzyme NADPH conversion cycle (30), fatty acid synthesis, fatty acid beta-oxidation, nitrogen catabolite repression, assimilation and scavenging, autophagy and protein turnover. Proteomics of isolated lipid droplets (31) identified over 250 lipid droplet-associated proteins including fatty acid synthesis genes, several putative lipases, a homolog of the lipolysis-regulating protein perilipin (32-34), vesicle trafficking proteins such as Rab GTPases and SNARE proteins, as well as several mitochondrial and peroxisomal proteins.

While these studies were unambiguous advances for the field, significant work remains to establish the genetic determinants of lipid accumulation in *R. toruloides*. Differential transcript or protein abundance under nitrogen limitation is suggestive of function in lipid accumulation, but transcriptional regulation and gene function are often poorly correlated in laboratory conditions (35). Similarly, sequestration in the lipid droplet may help regulate availability of some proteins for functions not necessarily related to lipid metabolism (36). More direct functional data would help the *R. toruloides* community prioritize this extensive list of genes for more detailed study and identify additional genes not identifiable by proteomic and transcriptomic methods. Finally, these studies highlighted dozens of genes with no known function, and hundreds more with only limited functional predictions. A more functional approach could shed more light on the most unique aspects of *R. toruloides* biology.

As the number of available fungal genomes has exploded in recent years (37, 38), tools to explore the function of uncharacterized genes have lagged behind sequencing capacity. High-throughput functional genomics approaches will help more effectively exploit our genomic resources. Fitness analysis on pooled mutant populations with DNA-based sequence barcodes (BarSeq) has proven to be a flexible, powerful approach for elucidating gene function in diverse species of bacteria (39-43) and some fungi (44). The combination of BarSeq with physical enrichment methods such as fluorescence-activated cell sorting has enabled genome-wide screens for phenotypes beyond simple growth (45). Early approaches required laborious construction of genome-wide libraries of targeted deletion mutants (38), but high-throughput sequencing has enabled random-insertion strategies in which barcoded transposons are mapped to insertion sites *en mass* (RB-TnSeq) (46, 47). Relatively low transformation efficiencies and a lack of functional transposon systems has been a limiting factor in the application of these techniques to diverse, non-model fungal species, however.

The plant-pathogenic bacteria *Agrobacterium tumefaciens* has evolved an efficient system to transfer virulence genes into eukaryotic cells. Once in the host cell, these transfer DNAs (T-DNAs) integrate randomly into the genome (46). *A. tumefaciens* mediated transformation (ATMT) has been used extensively to introduce exogenous DNA into plants (47-51), and has been demonstrated to transform diverse fungi at high efficiency (52). Recently, Esher et al. used ATMT followed by mutant selection and high-throughput sequencing to identify several mutants with altered cell wall biosynthesis in the human pathogen *Cryptococcus neoformans* (53). Their method is only viable for characterization of a small pool of highly enriched mutants, however.

In this study, we demonstrate the application of ATMT to the construction of a randomly barcoded, random insertion library (RB-TDNAseq) in *R. toruloides*. In mapping genomic locations of random insertions, we report the first full genome survey of essential genes in a basidiomycete fungi, consisting of 1337 probable essential genes including 36 genes unique to basidiomycetes. We demonstrate that our barcoded mutant library is an effective tool to rapidly assess mutant phenotypes by exploring fatty acid catabolism in *R. toruloides*, confirming that mitochondrial beta-oxidation is essential for fatty acid utilization in this species. We also show that some members of its expanded complement of perosixomal acyl-CoA dehydrogenases are necessary for growth on different fatty acids, suggesting substrate specificity or conditional optimality for each enzyme. We investigate perturbed lipid accumulation in the mutant pool by fractionation of the population by buoyancy and fluorescence activated cell sorting. We identify 150 genes with significant roles in lipid accumulation, notably genes involved in signaling cascades (28 genes), gene expression (15 genes), protein modification or trafficking (15 genes), ubiquitination or proteolysis (9 genes), autophagy (9 genes), and amino acid synthesis (8 genes). We also find evidence that tRNA modification effects lipid accumulation in *R. toruloides*, identifying 5 genes with likely roles in thiolation of tRNA wobble residues. These results significantly advance our understanding of lipid metabolism in *R toruloides*; identify key biological processes that should be explored and optimized in any oleaginous yeast engineered for lipid production; support emerging evidence of deep connections between lipid droplet dynamics, vesicular trafficking, and protein sorting; and demonstrate a general approach for barcoded mutagenesis that should enable functional genomics in a wide variety of fungal species.

## Results

### A functional genomics platform for *R. toruloides*

To enable functional genomics in *R. toruloides* IFO 0880, we first improved the existing genome assembly and annotation (36) using a combination of long-read PacBio sequencing for a more complete *de novo* assembly, a more comprehensive informatics approach for gene model predictions and functional annotation, and manual refinement of those models using evidence from mRNA sequencing (Genbank accession LCTV02000000, also available at the Mycocosm genome portal (38)), (see supplementary text for details). Summary tables of gene IDs, predicted functions, and probable orthologs in other systems are included in Supplementary file 1. For brevity, we will refer to *R. toruloides* genes by the common name for their *Saccharomyces cerevisiae* orthologs (e.g. *MET2*) when such orthologous relationships are unambiguous. Otherwise, we will give the Mycocosm protein ID, e.g. *RTO4_12154* and *RTO4_14576* are both orthologs of *GPD1*.

Because no method existed for high-throughput genetics in *R. toruloides*, we adapted established protocols for mapping barcoded transposon insertions (RB-TnSeq) (54), to mapping barcoded T-DNA insertions introduced with *Agrobacterium tumefaciens* mediated transformation (ATMT). We call this method RB-TDNAseq (Figure 1A). In brief, we generated a diverse library of binary ATMT plasmids bearing nourseothricin resistance cassettes with ~10 million unique 20 base-pair sequence ‘barcodes’ by efficient type IIs restriction enzyme cloning (55), introduced the library into *A. tumefaciens* EHA105 by electroporation, then transformed *R. toruloides* with ATMT. Using a TnSeq-like protocol, we mapped the unique locations of 293,613 individual barcoded T-DNA insertions in the *R. toruloides* genome (see supplementary text for details). Once insertion sites were associated with their barcodes, pooled fitness experiments were performed using a simple, scalable BarSeq protocol as previously described (56).

**Figure 1.**
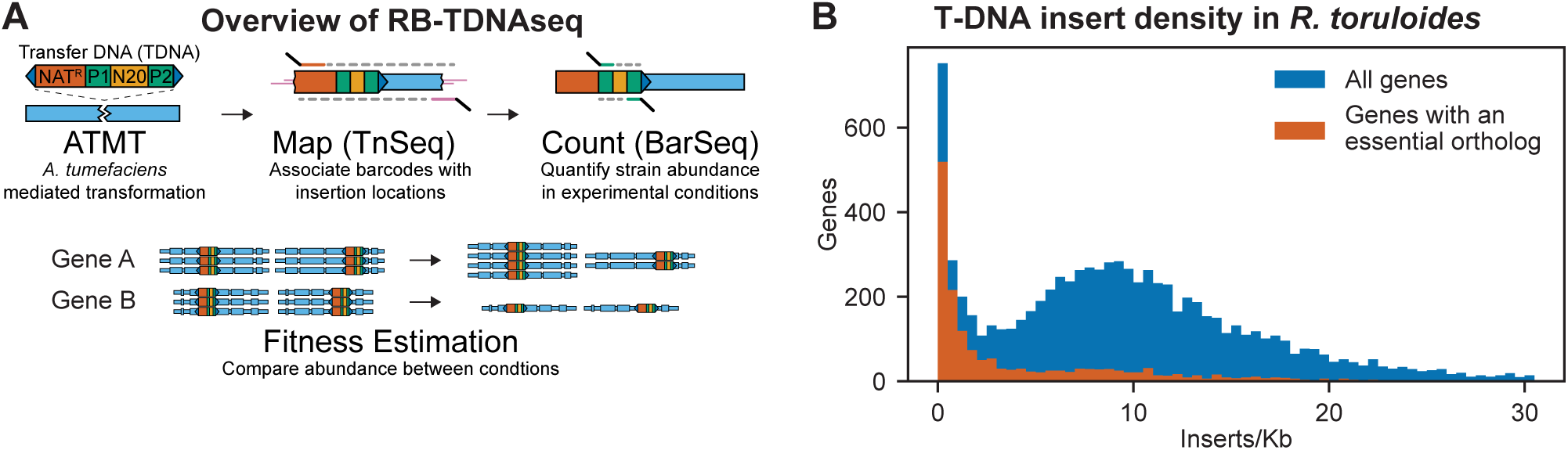
Overview of RB-TDNAseq and T-DNA insert density in *R. toruloides* coding regions. (A) General strategy of RB-TDNAseq. A library of binary plasmids bearing an antibiotic resistance cassette (NAT^R^) and a random 20 base-pair sequence ‘barcode’ (N20) flanked by specific priming sites (P1/P2) is introduced into a population of *A. tumefaciens* carrying a *vir* helper plasmid. *A. tumefaciens* efficiently transforms a T-DNA fragment into the target fungus (ATMT). NAT^R^ colonies are then combined to make a mutant pool. T-DNA-genome junctions are sequenced by TnSeq, thereby associating barcodes with the location of the insertion (Map). The mutant pool is then cultured under specific conditions and the relative abundance of mutant strains is measured by sequencing a short, specific, PCR on the barcodes (BarSeq) and counting the occurrence of each sequence (Count). Finally, for each gene, count data is combined across all barcodes mapping to insertions in that gene to obtain a robust measure of relative fitness for strains bearing mutations in that gene (Fitness Estimation). (B) Histogram of insert density in coding regions (start codon to stop codon) for all genes, and genes with orthologs reported to be essential in *A. nidulans, C. neoformans, N. crassa, S. cerevisiae*, or *S. pombe*.

Insertions were sufficiently well dispersed to map at least one T-DNA in 93% of nuclear genes, despite some local and fine-scale biases in insertion rates (see supplementary text for details). Insertion density in coding regions was consistently around 10 inserts/kb for most genes (Figure 1B). A subpopulation of genes with fewer than 2 inserts/kb was highly enriched for orthologs of genes reported as essential in *Aspergillus nidulans* (57), *Cryptococcus neoformans* (58), *Saccharomyces cerevisiae* (59), or *Schizosaccharomyces pombe* (60), or for which only heterokaryons could be obtained in the *Neurospora crassa* deletion collection (38). We therefore infer that these genes are essential in our library construction conditions, or at least that mutants for these genes have severely compromised growth. Based on the above criterion, we identified 1337 probable essential genes, which we report in Supplementary file 1. This list includes over 400 genes not reported as essential in the above-mentioned model fungi and is enriched for genes with homologs implicated in mitochondrial respiratory chain I assembly and function, dynein complex, the Swr1 complex, and mRNA non-sense mediated decay. For a full list of GO term enrichments see Supplementary file 1. This list also includes 36 genes unique to basidiomycetes.

### Mapping biosynthetic pathways using RB-TDNAseq

Before investigating more novel aspects of *R. toruloides’* biology, and to validate our methods, we wondered if RB-TDNAseq could be used to correctly annotate gene function in well-conserved biosynthetic pathways. Therefore, we investigated amino acid biosynthesis in *R. toruloides*. We cultured the mutant pool in defined medium (DM), consisting of yeast nitrogen base (YNB) and glucose, and in DM supplemented with amino acid and vitamins using drop-out complete mix (DOC). We then quantified insertion abundance in the starting and final populations using BarSeq. We applied the algorithms of Wetmore et al. (61) to compute an average fitness score (F) and T-like test statistic (T) for each gene. (see supplementary text for details) All fitness scores (averaged across biological replicates) and statistical tests reported here are available in Supplementary file 2 and online in a dynamic fitness browser, adapted from (62): http://fungalfit.genomics.lbl.gov/

Fitness scores for 6,558 genes in supplemented and non-supplemented media are shown in Figure 2A. Twenty-eight genes had significant, specific fitness defects in nons-upplemented media (T_DM-DOC_ < -3, F_DM_ < -1). Using an alternative, more conservative statistical approach (the Wilcoxon signed rank test (63, 64)), 23 of those genes were significantly less fit in non-supplemented media with a false discovery rate of 10% after multiple hypothesis correction (Supplementary file 2). When we grew the mutant pool in defined media with methionine or arginine supplementation (Figure 2B), these 28 genes for which mutants are auxotrophic partitioned into 11 mutants rescued by methionine, 10 mutants rescued by arginine, six mutants rescued by neither amino acid and one mutant rescued by both amino acids. All of the identified methionine and arginine auxotrophic mutants have orthologous genes for which mutants are auxotrophic for methionine/cysteine or arginine, respectively, in S. *cerevisiae* or *A. nidulans*. These data show that using RB-TDNAseq, we can repeatably identify critical genes for robust growth in an experimental condition and that an appropriate threshold for statistically and biologically significant fitness scores is |T| > 3. Of 31 genes involved in methionine and arginine metabolism that we could expect to identify with RB-TDNAseq, 30 met these statistical thresholds (see supplementary text). These data demonstrate that our BarSeq analysis should identify most, if not all, non-essential genes that are required for a specific biological process.

**Figure 2.**
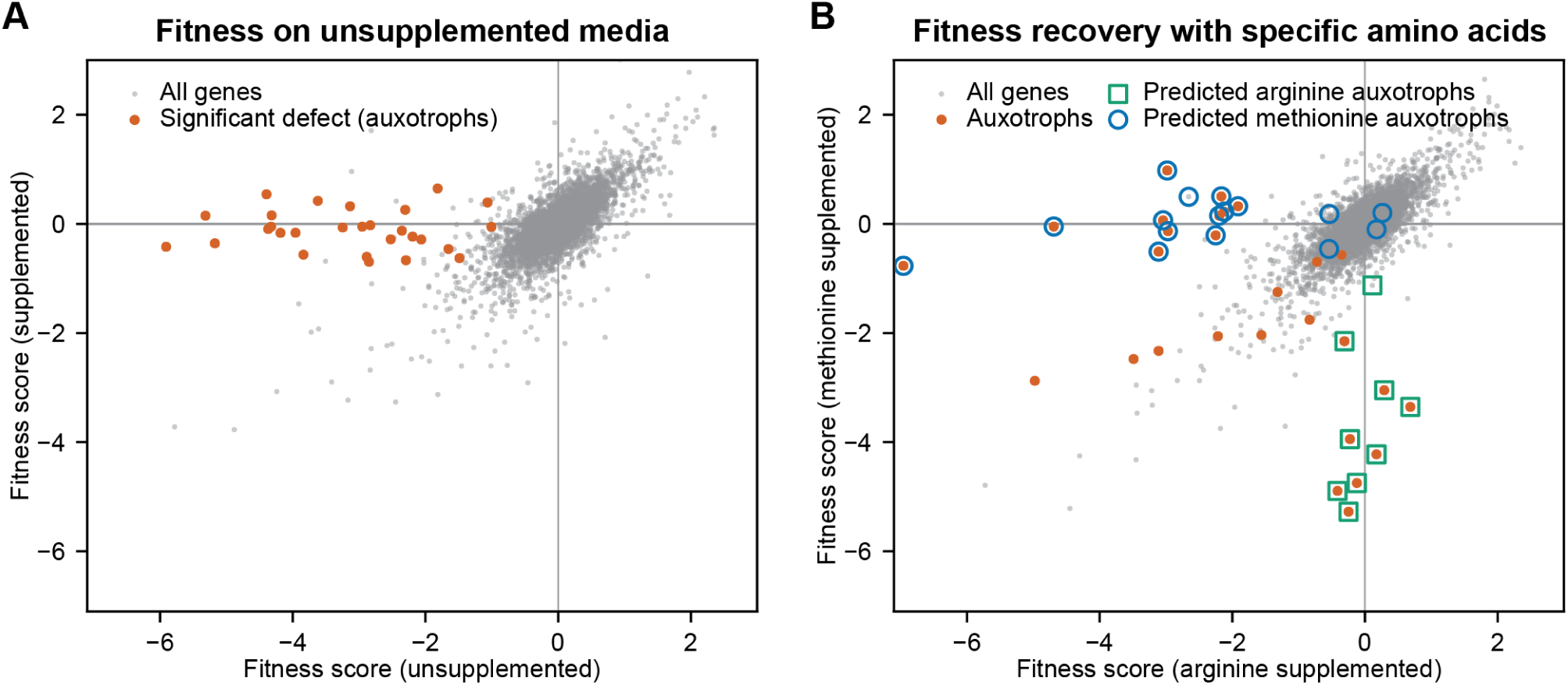
Confirmation of amino acid biosynthetic genes with high-throughput fitness experiments. (A) Fitness scores for 6,558 genes in media with and without amino acid supplementation (drop-out complete mix). Gene fitness scores are log ratios of final versus starting abundance averaged over multiple barcoded insertions per gene across 3 biological replicates. Genes that had significantly different enrichment scores between treatments (ΔF > 1, |T| statistic > 3) are highlighted and represent genes for which mutant strains are auxotrophic for one or more amino acids, nucleotides, or vitamins present in the drop-outcomplete mixture. (B) Fitness scores in media supplemented with arginine or methionine. Highlighted genes are the same as highlighted in (A). Deletion strains for circled or boxed genes are auxotrophic for methionine or arginine, respectively, in S. *cerevisiae* or *A. nidulans*. See supplementary file 2 for full fitness data.

### Fatty acid catabolism in *R. toruloides*

We next sought to understand how *R. toruloides* breaks down distinct fatty acids when used as growth substrates, as a window onto the complexity of lipid metabolism in this fungus. For this purpose, we used RB-TDNAseq to measure mutant fitness on three fatty acids as the sole carbon source: oleic acid (the most abundant fatty acid in *R. toruloides* (65, 66)), ricinoleic acid (a high-value fatty acid produced naturally in plants (67, 68) and synthetically in fungi (69)), and methyl-ricinoleic acid (a ricinoleic acid derivative used in lactone production (27, 70)). A total of 129 genes had significant fitness scores on one or more fatty acids including genes implicated in beta-oxidation of fatty acids, gluconeogenesis, mitochondrial amino acid metabolism, and several other aspects of cellular metabolism and gene regulation (See Figure 3 – Supplement 1 and the Supplemental text for a clustering analysis of fitness scores for these genes and Supplemental file 2 for a complete list).

We were particularly interested in beta-oxidation of fatty acids in the peroxisome and mitochondria, as these pathways are critical for lipid homeostasis (71-73), with major implications for both human health (74, 75) and metabolic engineering in fungi (76, 77). Fitness scores for *R. toruloides* genes homologous to enzymes with known roles in beta-oxidation of fatty acids are shown in Figure 3A. The localization for these enzymes is inferred mostly from homology to distantly related proteins in ascomycete fungi or mammalian species, but orthologs of five enzymes were localized to the predicted compartments using GFP fusion constructs in *Ustilago maydis* (78), demonstrating that localization is conserved across different species of basidiomycete fungi for at least some of these enzymes.

**Figure 3.**
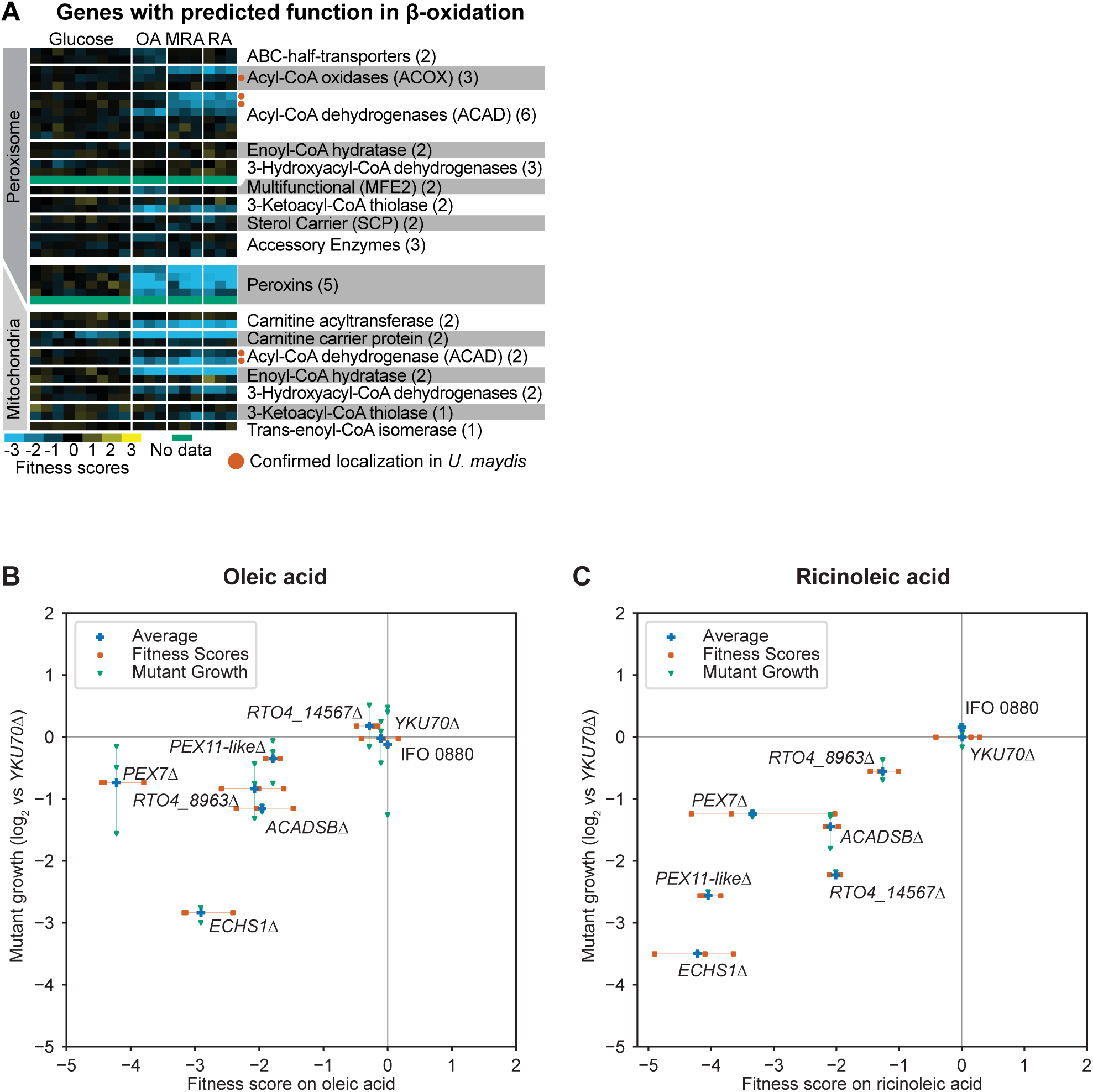
Genes with fitness defects on fatty acids. (A) Heatmap of fitness scores for *R. toruloides* genes with predicted roles in beta-oxidation of fatty acids. Enzyme classes and predicted locations were inferred from homologous proteins in *Ustilago maydis* as reported by Camoes et al (69). See supplementary file 2 for full fitness data. (B) Log_2_ optical density ratio for single deletion mutants versus the *YKU70*Δ control strain at mid-log phase on 1% oleic acid as carbon source are plotted against the fitness scores for each gene from BarSeq experiments on 1% oleic acid. (C) Log_2_ optical density ratio for single deletion mutants versus the *YKU70*Δ control strain at mid-log phase on 1% ricinoleic acid as carbon source are plotted against the fitness scores for mutants in each gene from BarSeq experiments on 1% ricinoleic acid.

Mutants for mitochondrial enzymes had the most consistent fitness scores across all three fatty acids, whereas mutants for the peroxisomal enzymes and peroxins had more variable fitness scores among fatty acids. Mutants for seven peroxisomal beta-oxidation enzymes and three peroxins had different fitness scores on oleic acid versus ricinoleic acid and methylricinoleic acid (listed in supplementary text, full fitness scores in Supplementary file 2), while 11 other predicted peroxisomal beta-oxidation enzymes had no significant fitness scores at all. The six predicted peroxisomal acyl-CoA dehydrogenases had particularly varied functional importance. Two were most important for ricinoleic acid and methylricinoelic acid utilization (*RTO4_10408* and *RTO4_14567*), one was most important for oleic acid utilization (*RTO4_8963*) and three were not necessary for growth on any of these fatty acids. These results demonstrate how RB-TDNAseq can be used to rapidly identify condition-specific functions among closely related members of a gene family for further genetic analysis or biochemical characterization. All together our data are consistent with a model of fatty acid beta-oxidation in *R. toruloides* in which diverse long-chain fatty acids are shortened in the peroxisome and a less structurally diverse set of short-chain fatty acids are oxidized to acetyl-CoA in the mitochondria (Figure 3 – figure supplement 2).

To validate our fitness data on fatty acids, we made targeted deletion mutants for several predicted peroxisomal and mitochondrial proteins. These strains were constructed by homologous recombination into a strain of *R. toruloides* IFO 0880 made deficient for non-homologous end joining by deleting *YKU70* (a.k.a. *KU70*) (14, 79, 80). We grew these mutant strains on oleic or ricinoleic acid media and compared their growth to the parental *YKU70*Δ strain in mid-log phase. Relative growth for the deletion strain for each gene is compared to its fitness scores in the BarSeq experiment in Figure 3B and Figure 3C. BarSeq fitness scores were reliable predictors of significant growth defects. The *PEX7*Δ mutant had similar fitness defects on both fatty acids, but mutants for *RTO4_8673* (similar to *PEX11*) and *RTO4_14567* (similar to *H. sapiens ACAD11*), had stronger fitness defects on ricinoleic acid, and the mutant for acyl-CoA dehydrogenase *RTO4_8963* had stronger fitness defects on oleic acid as predicted from fitness scores. Over a 96-hour time course, the *RTO4_14567*Δ mutant failed to grow at all on ricinoleic acid, whereas the *RTO4_8963*Δ mutant and the *PEX11* homolog *RTO4_8673*Δ mutant had more subtle phenotypes, approaching the same final density of the *YKU70*Δ control strain after a longer growth phase (Figure 3 – figure supplement 3).

### Functional Genomics of Lipid Accumulation in *R. toruloides*

To dissect the genetic basis of lipid accumulation in *R. toruloides*, we needed to extend our BarSeq methods beyond growth-based fitness assays. We induced lipid accumulation by nitrogen limitation (*R. toruloides* lipid droplets visualized in Figure 4A), and used two measures of cellular lipid content to fractionate the mutant pool (Figure 4B and supplementary text). We used the neutral-lipid stain BODIPY 493/503 (12) and fluorescence activated cell sorting (FACS) to enrich populations with larger/more or smaller/fewer lipid droplets (81). We also used buoyancy separation on sucrose gradients to enrich for populations with higher or lower total lipid content (41). Because many mutations can affect cell buoyant density independent of lipid accumulation (41), we also grew the mutant pool in rich media (YPD) and subjected it to sucrose gradient separation as a control for lipid-independent buoyancy phenotypes. For each pair of high and low lipid fractions, we then calculated an “enrichment score”, E, and T-statistic for each gene. E is analogous to our fitness scores based on growth, except that it is the log_2_ ratio of abundance in the high lipid fraction to the low lipid fraction, whereas F is the log_2_ ratio of final to initial abundance. Hierarchical clusters of enrichment scores for 271 genes for which mutants have significantly altered lipid accumulation (IEI > 1 and ITI > 3) are shown in Figure 5A. Enrichment scores, T-statistics, and multiple-hypothesis-adjusted significance scores from Wilcoxon signed rank tests for all 6,558 genes with sufficient BarSeq data are reported in Supplementary file 2.

**Figure 4.**
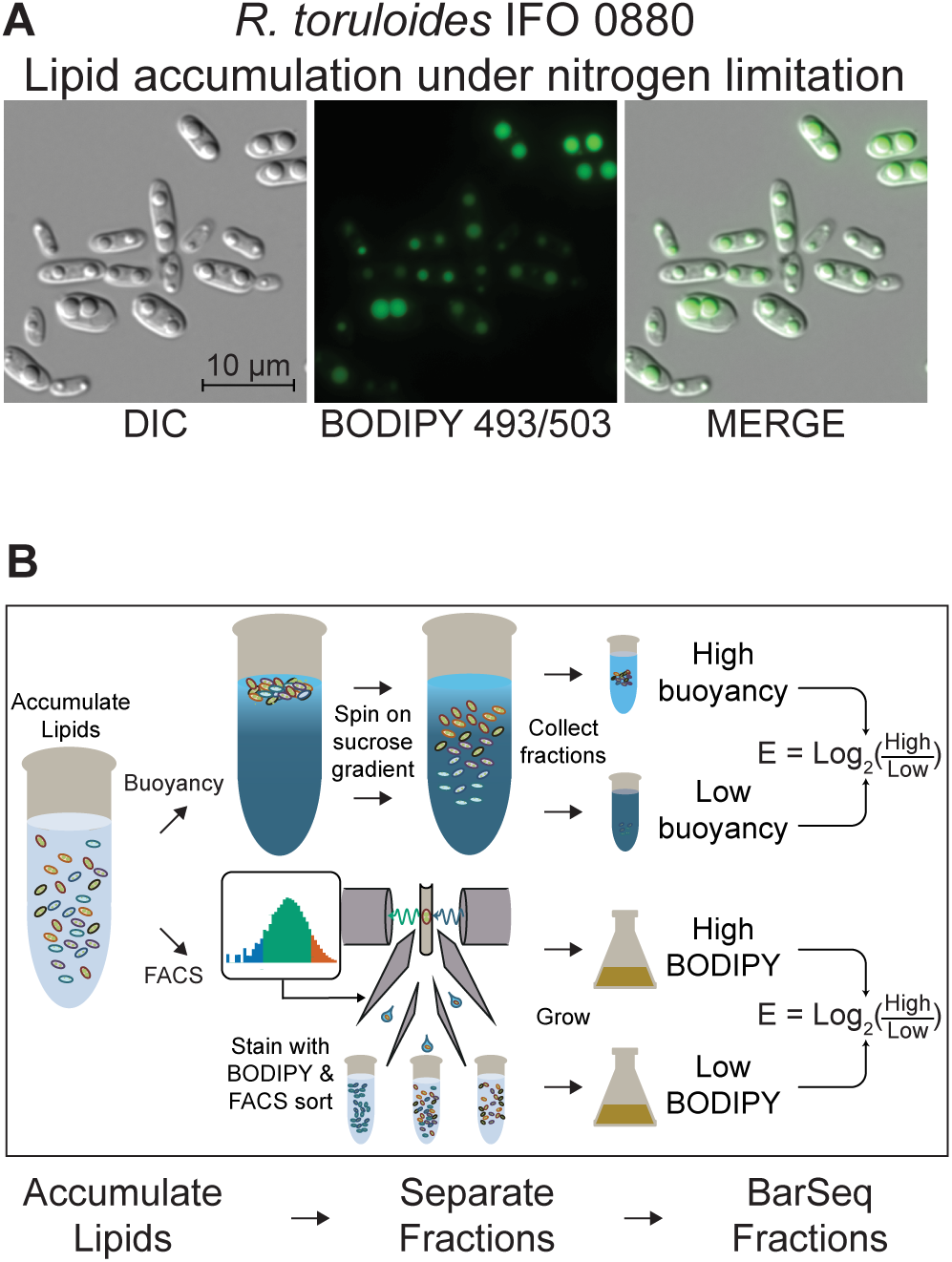
Detecting mutants with altered lipid accumulation. (A) Lipid accumulation in *R. toruloides* under nitrogen limitation. DIC microscopy of *R. toruloides* grown in low nitrogen media for 40 hours and stained with BODIPY 493/503 to label lipid droplets. (B) Two strategies to enrich populations for high or low TAG content cells. (Top) Buoyant density separation on sucrose gradients. Lipid accumulated cells are loaded on to a linear sucrose gradient and centrifuged. Cells settle at their neutral buoyancy, with the size of the low-density lipid droplet as the main driver of buoyancy differences. The gradient is then split into several fractions, and fractions representing the most and least buoyant 5-10% of the population, as well as a no-separation control are subjected to DNA extraction and strain quantification with BarSeq. For each gene an enrichment score is calculated as the log ratio of mutant abundance in the high buoyancy versus low buoyancy fractions. (Bottom) FACS sorting on BODIPY signal. Cells cultured in lipid accumulation conditions (limited nitrogen) are stained with BODIPY 493/503, then sorted in a FACS system. The 10% of the population with the highest and lowest BODIPY signal are sorted into enriched populations, as well as non-gated control. These small populations (10 million cells each) are then cultured for additional biomass and subjected to DNA extraction and strain quantification with BarSeq. For each gene, a FACS enrichment score is calculated as the log ratio of mutant abundance in the high BODIPY versus low BODIPY fractions.

**Figure 5.**
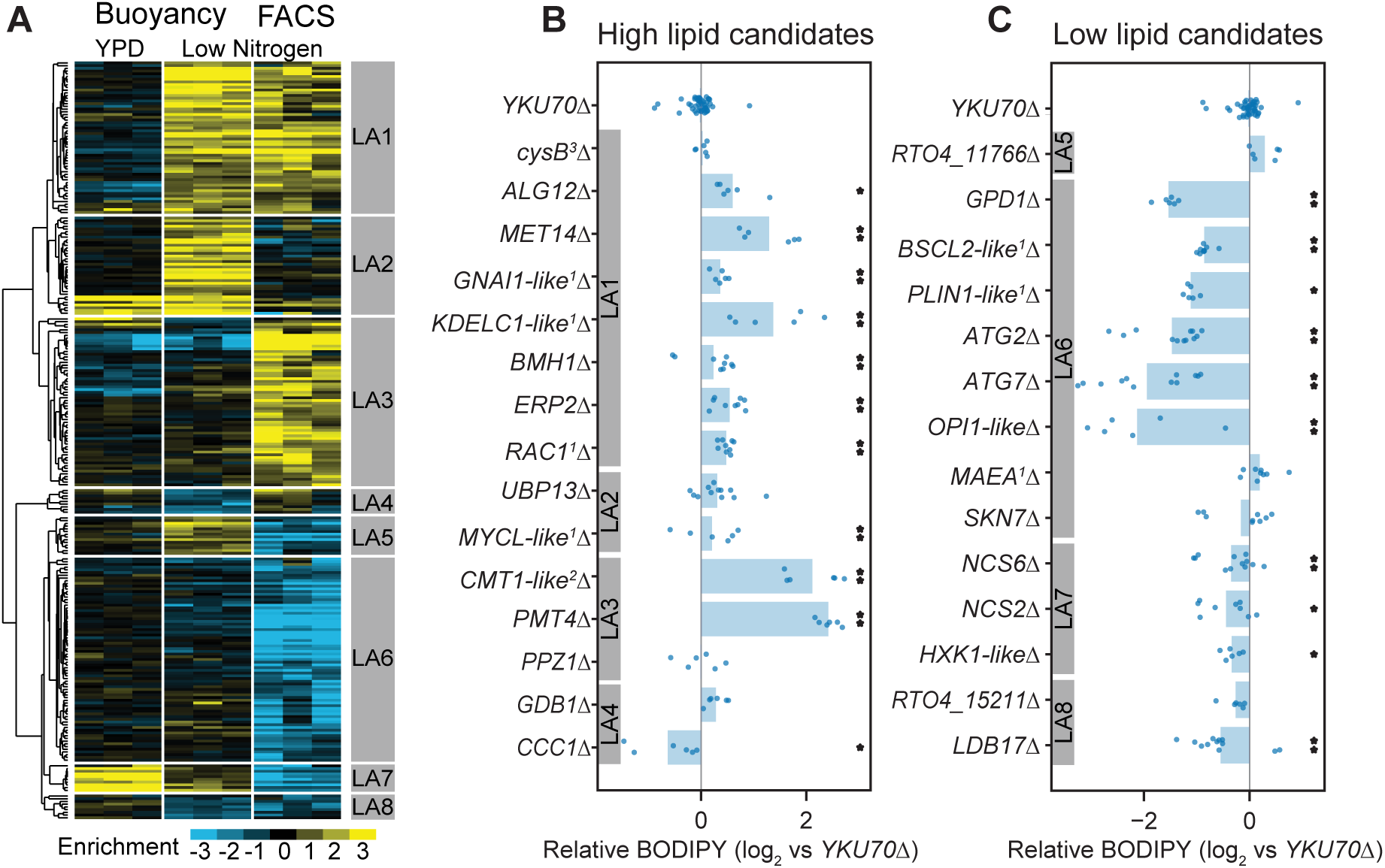
RB-TDNAseq on enriched populations identifies genes affecting lipid accumulation. (A) Hierarchical clusters of enrichment scores for 271 genes with significant enrichment (|E| > 1, |T| > 3) in high/low fractions separated by the buoyant density or FACS sorting of BODIPY stained cells after lipid accumulation on low nitrogen media. Enrichment scores for individual biological replicates (3 per condition) were clustered in this analysis. Eight major clusters were identified (LA1-LA8). See supplementary file 2 for full enrichment data. (B and C) Relative BODIPY signal for deletion mutants. Points are the average BODIPY/cell for 10,000 cells from independent biological replicate cultures normalized to three control *YKU70*Δ cultures processed on the same day. Three biological replicates were processed for each strain in any given experiment and each strain was included in at least two experiments processed on different days. ** P < 0.01, * P < 0.05 by one-tailed homoscedastic T-test versus *YKU70*Δ. ^1^Human homolog, ^2^*C. neoformans* homolog, ^3^*A. nidulans* homolog.

To assess the reliability of these enrichment scores in predicting phenotypes for null mutants, we constructed 29 single gene deletion mutants by homologous recombination in a *YKU70*Δ strain of IFO 0880 and measured lipid accumulation by average BODIPY fluorescence for 10,000 cells from each strain using flow cytometry (Figures 5B and 5C). When enrichment scores from both assays were strongly positive (LA1), we found that 7 of 8 deletion mutants had the expected phenotype (i.e. increased lipid accumulation). When only one assay yielded a strongly positive score (clusters LA2 and LA3), only 3 of 5 mutants had apparent increases in lipid content as measured by flow cytometry. Further, for the two mutants for genes in cluster LA3 with the greatest apparent increase in lipid content (*PMT4* and *RTO4_10302*, similar to *C. neoformans CMT1*) that measurement was likely an artifact of incomplete cell separation. Both mutants formed long chains of cells (see Figure 7 – figure supplement 1 for microscopy images), which would be analyzed as a single cell by our FACS assay. Genes in clusters LA4 and LA5 had conflicting enrichment scores between the two assays. Of three targeted deletion strains for genes in these clusters, only one (*CCC1*Δ) had a statistically significant phenotype, with decreased lipid accumulation. When the FACS assay gave a strongly negative score and there was no strong contrary buoyancy score (clusters LA6, LA7, and LA8), 10 of 13 mutants had reduced lipid accumulation. These data confirm that the both separation techniques are fundamentally sound, though in isolation each method has a significant rate of false positives. In combination, the two assays identified a large set of high-confidence candidate genes with important roles in lipid accumulation.

### Diverse predicted functions for lipid accumulation mutants

We manually curated homology-based predicted functions for the 393 genes with significant fitness or enrichment scores in this study (Supplementary file 1). An overview of predicted localization and functions for genes we identified with roles in fatty acid utilization or lipid accumulation is shown in Figure 6, with more detail for mutants with increased and decreased lipid accumulation in Tables 1 and 2, respectively. Note that we have excluded genes for which only one enrichment technique indicated altered lipid accumulation from this analysis. Mutants with increased lipid accumulation (cluster LA1, 56 genes) were most notably enriched for genes involved in signaling cascades, posttranslational protein modification and trafficking, and in amino acid biosynthesis. Mutants with decreased lipid accumulation (clusters LA6, LA7, and LA8, 94 genes) were most notably enriched for genes with roles in tRNA-modification; regulatory kinases and phosphatases; and genes involved in cellular recycling processes such as autophagy, ubiquitin-protease systems and the unfolded protein response.

**Figure 6.**
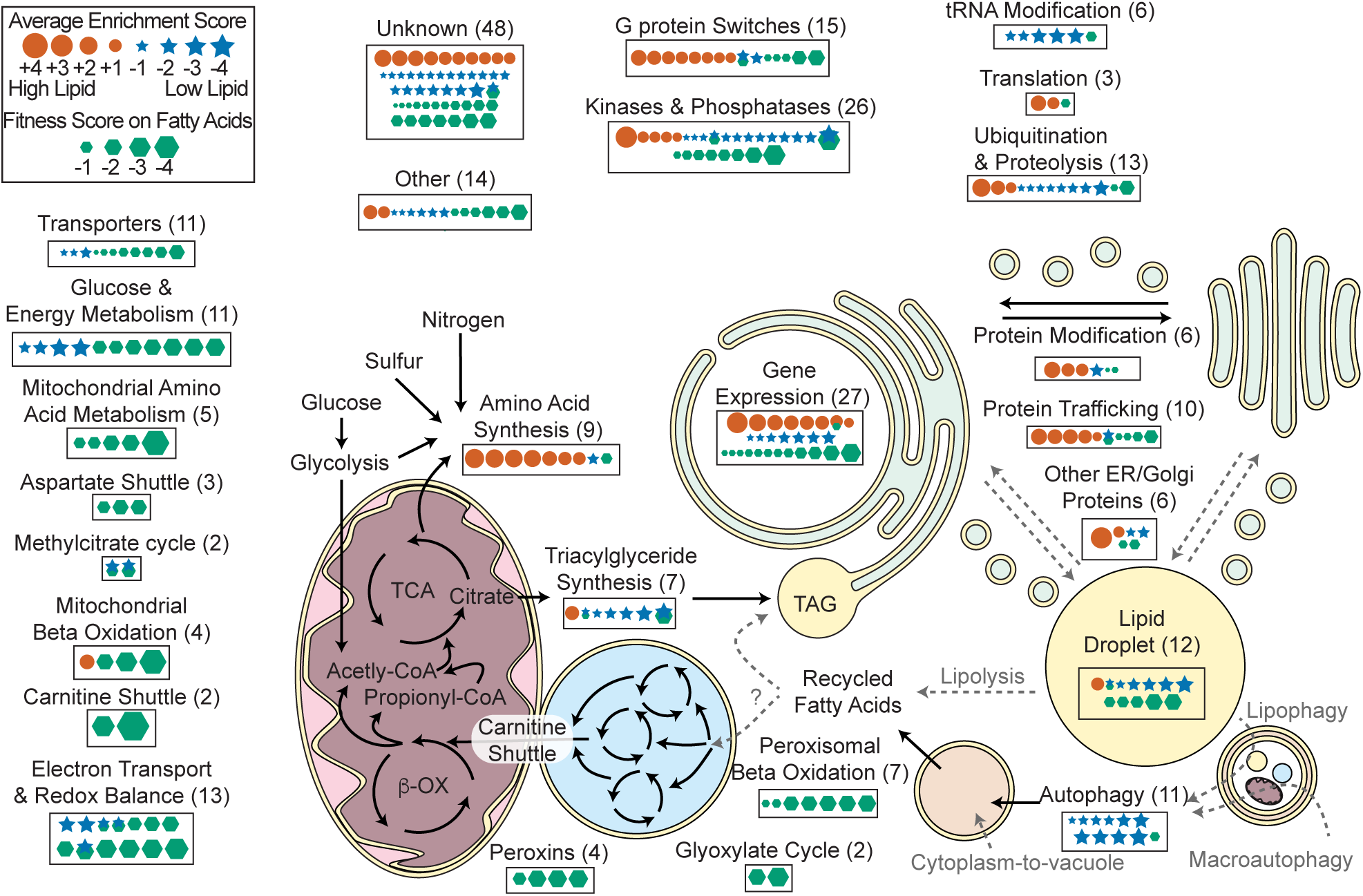
Overview of *R. toruloides* lipid metabolism. Key metabolic pathways and cellular functions mediating lipid metabolism as identified from fitness scores on fatty acid and enrichment scores from lipid accumulation screens. Fitness and/or enrichment scores for individual genes are depicted graphically by relative size of hexagonal, circular or star icons respectively. Only fitness scores for genes with significant growth defects on at least one fatty acid (see supplementary file 2) and enrichment scores from high confidence clusters (see Figure 5 and supplementary file 2) are shown. Enrichment scores were averaged between buoyancy and FACS experiments, except for genes with confounding enrichment scores in rich media conditions, for which only FACS data were averaged. Positive scores (orange circles) represent genes for which mutants have increased lipid accumulation. Negative fitness scores (blue stars) represent genes for which mutants have decreased lipid accumulation. Genes detected in proteomics of *R. toruloides* lipid droplets by Zhu et al (*RAC1, GUT2, PLIN1, EGH1, RIP1, MGL2, AAT1, CIR2, MLS1*, and *RTO4_8963*) or found in lipid droplet of many organisms (*DGA1* and *BSCL2*)(see Supplementary File 5) are depicted under “Lipid Droplet” and also their molecular functions, e.g. “G Protein Switches” for *RAC1*.

**Table 1.**
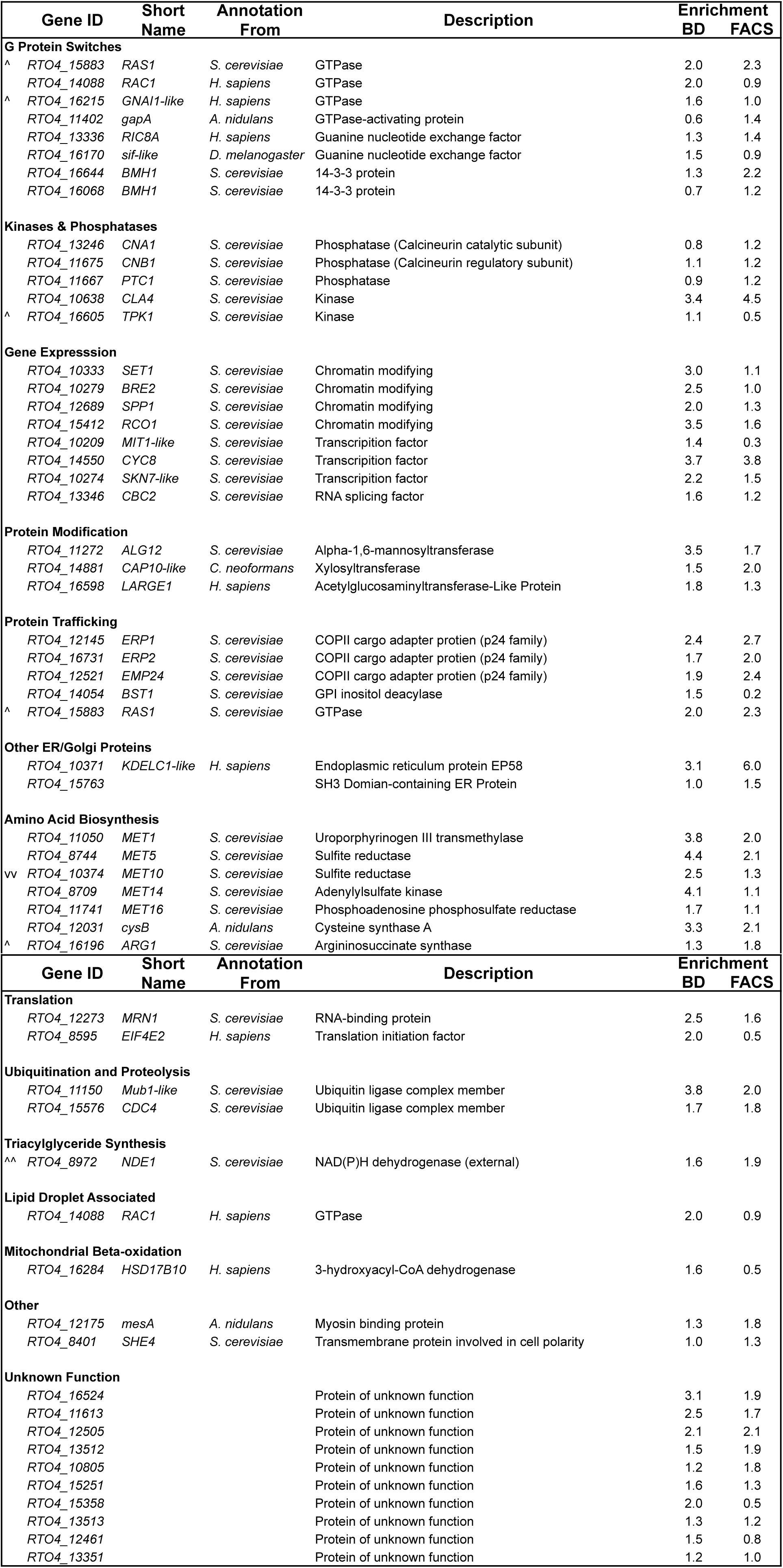
Predicted gene function: Mutants with increased lipid accumulation. Predicted functions for genes for which mutants were high-confidence candidates for increased lipid accumulation (enrichment scores clustered in LA1, Figure 5). Cellular processes grouped as in Figure 6. BD: Enrichment score from buoyant density separation. FACS: Enrichment score from fluorescence activated cell sorting. ^^^ Protein abundance increased under nitrogen limitation Zhu et al 2012 (8). ^^^^ Protein abundance increase 10-fold or more. v Protein abundance decreased. vv Protein abundance decreased 10-fold or more.

**Table 2.**
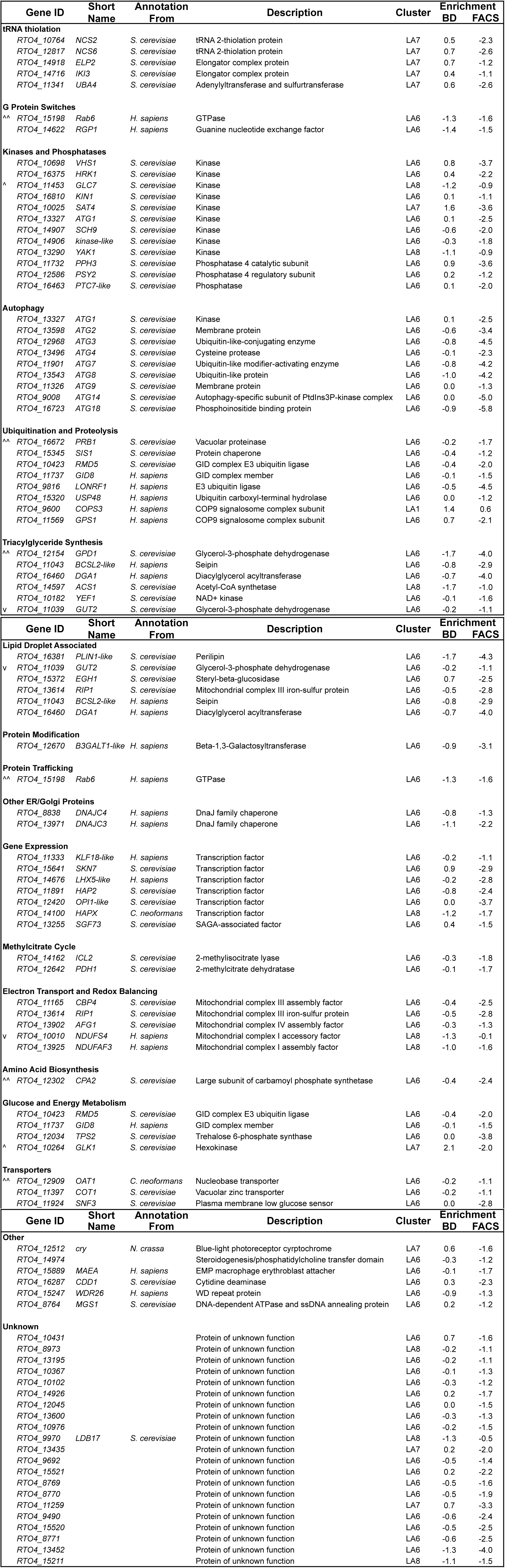
Predicted gene function: Mutants with decreased lipid accumulation. Predicted functions for genes for which mutants were high-confidence candidates for decreased lipid accumulation (enrichment scores clustered in LA6 - LA8, Figure 5). Cellular processes grouped as in Figure 6. BD: Enrichment score from buoyant density separation. FACS: Enrichment score from fluorescence activated cell sorting. ^^^ Protein abundance increased under nitrogen limitation Zhu et al 2012 (8). ^^^^ Protein abundance increase 10-fold or more. v Protein abundance decreased. vv Protein abundance decreased 10-fold or more.

| Gene ID | Short Name | Annotation From | Description | Cluster | Enrichment BD | FACS |
| --- | --- | --- | --- | --- | --- | --- |
| **tRNA thiolation** |  |  |  |  |  |  |
| *RTO4_10764* | *NCS2* | *S. cerevisiae* | tRNA 2-thiolation protein | LA7 | 0.5 | -2.3 |
| *RTO4_12817* | *NCS6* | *S. cerevisiae* | tRNA 2-thiolation protein | LA7 | 0.7 | -2.6 |
| *RTO4_14918* | *ELP2* | *S. cerevisiae* | Elongator complex protein | LA7 | 0.7 | -1.2 |
| *RTO4_14716* | *IKI3* | *S. cerevisiae* | Elongator complex protein | LA7 | 0.4 | -1.1 |
| *RTO4_11341* | *UBA4* | *S. cerevisiae* | Adenylyltransferase and sulfurtransferase | LA7 | 0.6 | -2.6 |
| **G Protein Switches** |  |  |  |  |  |  |
| <sup>^^</sup> *RTO4_15198* | *Rab6* | *H. sapiens* | GTPase | LA6 | -1.3 | -1.6 |
| *RTO4_14622* | *RGP1* | *H. sapiens* | Guanine nucleotide exchange factor | LA6 | -1.4 | -1.5 |
| **Kinases and Phosphatases** |  |  |  |  |  |  |
| *RTO4_10698* | *VHS1* | *S. cerevisiae* | Kinase | LA6 | 0.8 | -3.7 |
| *RTO4_16375* | *HRK1* | *S. cerevisiae* | Kinase | LA6 | 0.4 | -2.2 |
| <sup>^</sup> *RTO4_11453* | *GLC7* | *S. cerevisiae* | Kinase | LA8 | -1.2 | -0.9 |
| *RTO4_16810* | *KIN1* | *S. cerevisiae* | Kinase | LA6 | 0.1 | -1.1 |
| *RTO4_10025* | *SAT4* | *S. cerevisiae* | Kinase | LA7 | 1.6 | -3.6 |
| *RTO4_13327* | *ATG1* | *S. cerevisiae* | Kinase | LA6 | 0.1 | -2.5 |
| *RTO4_14907* | *SCH9* | *S. cerevisiae* | Kinase | LA6 | -0.6 | -2.0 |
| *RTO4_14906* | *kinase-like* | *S. cerevisiae* | Kinase | LA6 | -0.3 | -1.8 |
| *RTO4_13290* | *YAK1* | *S. cerevisiae* | Kinase | LA8 | -1.1 | -0.9 |
| *RTO4_11732* | *PPH3* | *S. cerevisiae* | Phosphatase 4 catalytic subunit | LA6 | 0.9 | -3.6 |
| *RTO4_12586* | *PSY2* | *S. cerevisiae* | Phosphatase 4 regulatory subunit | LA6 | 0.2 | -1.2 |
| *RTO4_16463* | *PTC7-like* | *S. cerevisiae* | Phosphatase | LA6 | 0.1 | -2.0 |
| **Autophagy** |  |  |  |  |  |  |
| *RTO4_13327* | *ATG1* | *S. cerevisiae* | Kinase | LA6 | 0.1 | -2.5 |
| *RTO4_13598* | *ATG2* | *S. cerevisiae* | Membrane protein | LA6 | -0.6 | -3.4 |
| *RTO4_12968* | *ATG3* | *S. cerevisiae* | Ubiquitin-like-conjugating enzyme | LA6 | -0.8 | -4.5 |
| *RTO4_13496* | *ATG4* | *S. cerevisiae* | Cysteine protease | LA6 | -0.1 | -2.3 |
| *RTO4_11901* | *ATG7* | *S. cerevisiae* | Ubiquitin-like modifier-activating enzyme | LA6 | -0.8 | -4.2 |
| *RTO4_13543* | *ATG8* | *S. cerevisiae* | Ubiquitin-like protein | LA6 | -1.0 | -4.2 |
| *RTO4_11326* | *ATG9* | *S. cerevisiae* | Membrane protein | LA6 | 0.0 | -1.3 |
| *RTO4_9008* | *ATG14* | *S. cerevisiae* | Autophagy-specific subunit of PtdIns3P-kinase complex | LA6 | 0.0 | -5.0 |
| *RTO4_16723* | *ATG18* | *S. cerevisiae* | Phosphoinositide binding protein | LA6 | -0.9 | -5.8 |
| **Ubiquitination and Proteolysis** |  |  |  |  |  |  |
| <sup>^^</sup> *RTO4_16672* | *PRB1* | *S. cerevisiae* | Vacuolar proteinase | LA6 | -0.2 | -1.7 |
| *RTO4_15345* | *SIS1* | *S. cerevisiae* | Protein chaperone | LA6 | -0.4 | -1.2 |
| *RTO4_10423* | *RMD5* | *S. cerevisiae* | GID complex E3 ubiquitin ligase | LA6 | -0.4 | -2.0 |
| *RTO4_11737* | *GID8* | *H. sapiens* | GID complex member | LA6 | -0.1 | -1.5 |
| *RTO4_9816* | *LONRF1* | *H. sapiens* | E3 ubiquitin ligase | LA6 | -0.5 | -4.5 |
| *RTO4_15320* | *USP48* | *H. sapiens* | Ubiquitin carboxyl-terminal hydrolase | LA6 | 0.0 | -1.2 |
| *RTO4_9600* | *COPS3* | *H. sapiens* | COP9 signalosome complex subunit | LA1 | 1.4 | 0.6 |
| *RTO4_11569* | *GPS1* | *H. sapiens* | COP9 signalosome complex subunit | LA6 | 0.7 | -2.1 |
| **Triacylglyceride Synthesis** |  |  |  |  |  |  |
| <sup>^^</sup> *RTO4_12154* | *GPD1* | *S. cerevisiae* | Glycerol-3-phosphate dehydrogenase | LA6 | -1.7 | -4.0 |
| *RTO4_11043* | *BCSL2-like* | *H. sapiens* | Seipin | LA6 | -0.8 | -2.9 |
| *RTO4_16460* | *DGA1* | *H. sapiens* | Diacylglycerol acyltransferase | LA6 | -0.7 | -4.0 |
| *RTO4_14597* | *ACS1* | *S. cerevisiae* | Acetyl-CoA synthetase | LA8 | -1.7 | -1.0 |
| *RTO4_10182* | *YEF1* | *S. cerevisiae* | NAD<sup>+</sup> kinase | LA6 | -0.1 | -1.6 |
| <sup>v</sup> *RTO4_11039* | *GUT2* | *S. cerevisiae* | Glycerol-3-phosphate dehydrogenase | LA6 | -0.2 | -1.1 |

**Table 2. Predicted gene function: Mutants with decreased lipid accumulation. (continued)**
| Gene ID | Short Name | Annotation From | Description | Cluster | Enrichment BD | FACS |
| --- | --- | --- | --- | --- | --- | --- |
| **Lipid Droplet Associated** |  |  |  |  |  |  |
| *RTO4_16381* | *PLIN1-like* | *S. cerevisiae* | Perilipin | LA6 | -1.7 | -4.3 |
| v *RTO4_11039* | *GUT2* | *S. cerevisiae* | Glycerol-3-phosphate dehydrogenase | LA6 | -0.2 | -1.1 |
| *RTO4_15372* | *EGH1* | *S. cerevisiae* | Steryl-beta-glucosidase | LA6 | 0.7 | -2.5 |
| *RTO4_13614* | *RIP1* | *S. cerevisiae* | Mitochondrial complex III iron-sulfur protein | LA6 | -0.5 | -2.8 |
| *RTO4_11043* | *BCSL2-like* | *H. sapiens* | Seipin | LA6 | -0.8 | -2.9 |
| *RTO4_16460* | *DGA1* | *H. sapiens* | Diacylglycerol acyltransferase | LA6 | -0.7 | -4.0 |
| **Protein Modification** |  |  |  |  |  |  |
| *RTO4_12670* | *B3GALT1-like* | *H. sapiens* | Beta-1,3-Galactosyltransferase | LA6 | -0.9 | -3.1 |
| **Protein Trafficking** |  |  |  |  |  |  |
| ^^ *RTO4_15198* | *Rab6* | *H. sapiens* | GTPase | LA6 | -1.3 | -1.6 |
| **Other ER/Golgi Proteins** |  |  |  |  |  |  |
| *RTO4_8838* | *DNAJC4* | *H. sapiens* | DnaJ family chaperone | LA6 | -0.8 | -1.3 |
| *RTO4_13971* | *DNAJC3* | *H. sapiens* | DnaJ family chaperone | LA6 | -1.1 | -2.2 |
| **Gene Expression** |  |  |  |  |  |  |
| *RTO4_11333* | *KLF18-like* | *H. sapiens* | Transcription factor | LA6 | -0.2 | -1.1 |
| *RTO4_15641* | *SKN7* | *S. cerevisiae* | Transcription factor | LA6 | 0.9 | -2.9 |
| *RTO4_14676* | *LHX5-like* | *H. sapiens* | Transcription factor | LA6 | -0.2 | -2.8 |
| *RTO4_11891* | *HAP2* | *S. cerevisiae* | Transcription factor | LA6 | -0.8 | -2.4 |
| *RTO4_12420* | *OPI1-like* | *S. cerevisiae* | Transcription factor | LA6 | 0.0 | -3.7 |
| *RTO4_14100* | *HAPX* | *C. neoformans* | Transcription factor | LA8 | -1.2 | -1.7 |
| *RTO4_13255* | *SGF73* | *S. cerevisiae* | SAGA-associated factor | LA6 | 0.4 | -1.5 |
| **Methylcitrate Cycle** |  |  |  |  |  |  |
| *RTO4_14162* | *ICL2* | *S. cerevisiae* | 2-methylisocitrate lyase | LA6 | -0.3 | -1.8 |
| *RTO4_12642* | *PDH1* | *S. cerevisiae* | 2-methylcitrate dehydratase | LA6 | -0.1 | -1.7 |
| **Electron Transport and Redox Balancing** |  |  |  |  |  |  |
| *RTO4_11165* | *CBP4* | *S. cerevisiae* | Mitochondrial complex III assembly factor | LA6 | -0.4 | -2.5 |
| *RTO4_13614* | *RIP1* | *S. cerevisiae* | Mitochondrial complex III iron-sulfur protein | LA6 | -0.5 | -2.8 |
| *RTO4_13902* | *AFG1* | *S. cerevisiae* | Mitochondrial complex IV assembly factor | LA6 | -0.3 | -1.3 |
| v *RTO4_10010* | *NDUFS4* | *H. sapiens* | Mitochondrial complex I accessory factor | LA8 | -1.3 | -0.1 |
| *RTO4_13925* | *NDUFAF3* | *H. sapiens* | Mitochondrial complex I assembly factor | LA8 | -1.0 | -1.6 |
| **Amino Acid Biosynthesis** |  |  |  |  |  |  |
| ^^ *RTO4_12302* | *CPA2* | *S. cerevisiae* | Large subunit of carbamoyl phosphate synthetase | LA6 | -0.4 | -2.4 |
| **Glucose and Energy Metabolism** |  |  |  |  |  |  |
| *RTO4_10423* | *RMD5* | *S. cerevisiae* | GID complex E3 ubiquitin ligase | LA6 | -0.4 | -2.0 |
| *RTO4_11737* | *GID8* | *H. sapiens* | GID complex member | LA6 | -0.1 | -1.5 |
| *RTO4_12034* | *TPS2* | *S. cerevisiae* | Trehalose 6-phosphate synthase | LA6 | 0.0 | -3.8 |
| ^ *RTO4_10264* | *GLK1* | *S. cerevisiae* | Hexokinase | LA7 | 2.1 | -2.0 |
| **Transporters** |  |  |  |  |  |  |
| ^^ *RTO4_12909* | *OAT1* | *C. neoformans* | Nucleobase transporter | LA6 | -0.2 | -1.1 |
| *RTO4_11397* | *COT1* | *S. cerevisiae* | Vacuolar zinc transporter | LA6 | -0.2 | -1.1 |
| *RTO4_11924* | *SNF3* | *S. cerevisiae* | Plasma membrane low glucose sensor | LA6 | 0.0 | -2.8 |

**Table 2. Predicted gene function: Mutants with decreased lipid accumulation. (continued)**
| Gene ID | Short Name | Annotation From | Description | Cluster | Enrichment BD | FACS |
| --- | --- | --- | --- | --- | --- | --- |
| **Other** |  |  |  |  |  |  |
| *RTO4_12512* | *cry* | *N. crassa* | Blue-light photoreceptor cyrptochrome | LA7 | 0.6 | -1.6 |
| *RTO4_14974* |  |  | Steroidogenesis/phosphatidylcholine transfer domain | LA6 | -0.3 | -1.2 |
| *RTO4_15889* | *MAEA* | *H. sapiens* | EMP macrophage erythroblast attacher | LA6 | -0.1 | -1.7 |
| *RTO4_16287* | *CDD1* | *S. cerevisiae* | Cytidine deaminase | LA6 | 0.3 | -2.3 |
| *RTO4_15247* | *WDR26* | *H. sapiens* | WD repeat protein | LA6 | -0.9 | -1.3 |
| *RTO4_8764* | *MGS1* | *S. cerevisiae* | DNA-dependent ATPase and ssDNA annealing protein | LA6 | 0.2 | -1.2 |
| **Unknown** |  |  |  |  |  |  |
| *RTO4_10431* |  |  | Protein of unknown function | LA6 | 0.7 | -1.6 |
| *RTO4_8973* |  |  | Protein of unknown function | LA8 | -0.2 | -1.1 |
| *RTO4_13195* |  |  | Protein of unknown function | LA6 | -0.2 | -1.1 |
| *RTO4_10367* |  |  | Protein of unknown function | LA6 | -0.1 | -1.3 |
| *RTO4_10102* |  |  | Protein of unknown function | LA6 | -0.3 | -1.2 |
| *RTO4_14926* |  |  | Protein of unknown function | LA6 | 0.2 | -1.7 |
| *RTO4_12045* |  |  | Protein of unknown function | LA6 | 0.0 | -1.5 |
| *RTO4_13600* |  |  | Protein of unknown function | LA6 | -0.3 | -1.3 |
| *RTO4_10976* |  |  | Protein of unknown function | LA6 | -0.2 | -1.5 |
| *RTO4_9970* | *LDB17* | *S. cerevisiae* | Protein of unknown function | LA8 | -1.3 | -0.5 |
| *RTO4_13435* |  |  | Protein of unknown function | LA7 | 0.2 | -2.0 |
| *RTO4_9692* |  |  | Protein of unknown function | LA6 | -0.5 | -1.4 |
| *RTO4_15521* |  |  | Protein of unknown function | LA6 | 0.2 | -2.2 |
| *RTO4_8769* |  |  | Protein of unknown function | LA6 | -0.5 | -1.6 |
| *RTO4_8770* |  |  | Protein of unknown function | LA6 | -0.5 | -1.9 |
| *RTO4_11259* |  |  | Protein of unknown function | LA7 | 0.7 | -3.3 |
| *RTO4_9490* |  |  | Protein of unknown function | LA6 | -0.6 | -2.4 |
| *RTO4_15520* |  |  | Protein of unknown function | LA6 | -0.5 | -2.5 |
| *RTO4_8771* |  |  | Protein of unknown function | LA6 | -0.6 | -2.5 |
| *RTO4_13452* |  |  | Protein of unknown function | LA6 | -1.3 | -4.0 |
| *RTO4_15211* |  |  | Protein of unknown function | LA8 | -1.1 | -1.5 |

#### Mutants with increased lipid accumulation

Mutants in several homologs of known signaling genes had increased lipid accumulation, depicted in Figure 6 under “G Protein Switches”, “Kinases & Phosphatases”, and “Gene Expression”. Three GTPases, a GTPase-activating protein (GAP) and two guanine nucleotide exchange factors (GEFs) were in cluster LA1, along with two orthologs of *BMH1*. BMH1 is a 14-3-3 family protein, involved in G protein signaling, the RAS/MAPK signaling cascade, and many other processes (82). The genes encoding calcineurin complex were also in this cluster as was another protein phosphatase and two protein kinases. Four genes with predicted roles in histone modification were included in cluster LA1 along with three transcription factors and the RNA splicing factor *CBC2*, which is involved in mRNA processing and degradation (83).

Mutants in ten genes with likely roles in protein modification, protein trafficking or other processes in the ER and Golgi led to increased lipid accumulation (Figure 6). These genes included three cargo adapter proteins, GPI anchor modifying protein *BST1*, the GTPase *RAS1* (which has been implicated in regulation of vesicular trafficking), and three probable glycosyltransferases. These results show that protein trafficking plays an important role in lipid accumulation in *R. toruloides*, as has been shown in other systems (84), though different ensembles of trafficking proteins may be involved in different species.

Disruption of sulfur assimilation also increased lipid accumulation, with five genes involved in sulfate conversion to sulfide clustering in LA1. The cysteine synthase *cysB* was also in this cluster, though *cysB*Δ mutants did not have significantly increased lipid accumulation in our flow cytometry assay. A *MET14*Δ mutant had significantly increased lipid content as expected (Figure 5B). In general, the sulfate assimilation mutants had reduced growth in low nitrogen conditions, as indicated by negative fitness scores for pre-enrichment control samples (Supplementary file 2). As expected, the auxotrophic mutants identified in our supplementation experiments also had compromised growth in low nitrogen conditions, though the phenotype was generally less severe, likely reflective of slower growth of the population generally. However, slower growth due to auxotrophy was not predictive of higher enrichment scores even for *MET2, MET6, MET12, and MET13*, which are required for methionine synthesis but not sulfate incorporation through cysteine (Figure 2 – figure supplement 2A). These data suggest that cysteine or intermediate sulfur compounds in the assimilation of sulfate to sulfide may be involved in regulation of lipid accumulation.

#### Mutants with decreased lipid accumulation

We found evidence that tRNA thiolation plays a role in lipid accumulation in *R. toruloides*. Enrichment scores for six genes known to be important in the thiolation of tRNA wobble residues (85) clustered together in LA7. Though these mutants also had apparent buoyancy phenotypes on YPD, two deletion strains (NCS6Δ and NCS2A) had reduced lipid content in pure culture (Figure 5C). Furthermore, we observed that for orthologs of *S. cerevisiae* genes with measured tRNA thiolation levels (86), a decrease in tRNA thiolation corresponded to a lower enrichment score (Figure 5 – figure supplement 1). Modification of tRNA wobble positions has been implicated in regulation of gene expression in response to heat shock (87) and sulfur (88) availability. Our observations suggest that in *R. toruloides* the refactoring of the proteome for efficient lipid accumulation requires fully functional tRNA thiolation. The role that tRNA thiolation plays in this metabolic transition is unclear and deserves more detailed study.

Efficient lipid accumulation also required the regulatory action of orthologs to the *H. sapiens* GTPase *Rab6* and the guanine nucleotide exchange factor *RGP1*, 9 protein kinases, 3 phosphatases or their binding partners. These genes are likely involved in signaling pathways mediating nutrient state. They include four genes with orthologs implicated in the regulation of glucose and glycogen metabolism (*VHS1, HRK1, GLC7* and *KIN1*) and four genes with orthologs involved in regulation of nitrogen catabolism (*PPH3, PSY2, SCH9*, and *ATG1*).

Mutants in nine core components of autophagy were deficient for lipid accumulation. The vacuolar protease *PRB1* and *SIS1* (chaperone mediating protein delivery to the proteasome) were also required for efficient lipid accumulation, as were six genes implicated in protein ubiquitination (Table 2). Ubiquitination can affect many aspects of gene function, but likely most of these genes participate in regulation of proteolysis. These results show that autophagy and recycling of cellular components are important for efficient lipid accumulation in *R. toruloides* and provide direct genetic evidence for a previous observation that chemical inhibition of autophagy using 3-methyladenine reduced lipid accumulation in the oleaginous yeast *Y. lipolytica* (89).

While most genes encoding enzymatic steps in fatty acid and TAG biosynthesis had too few insertions to calculate reliable enrichment scores (many are probable essential genes, see Supplementary file 1), mutants in six genes with predicted function in TAG synthesis resulted in lower lipid accumulation (see Figure 6 – figure supplement 1). Three of these genes directly mediated reactions in TAG synthesis: *RTO4_12154, RTO4_11043, and DGA1*. *RTO4_12154* is one of two *R. toruloides GPD1* orthologs predicted to convert dihydroxyacetone phosphate (DHAP) into glycerol-3-phosphate (G3P) (85). *RTO4_11043* is a distant homolog of *H. sapiens BSCL2* (seipin), which modulates the activity of G3P acyltransferase in nascent lipid droplets (90). *DGA1* catalyzes conversion of diacylglyceride into TAG (67). Three more genes were more peripherally involved in TAG biosynthesis: *ACS1, YEF1, and GUT2*. *ACS1*, acetyl-CoA synthetase (31), may supplement production of cytosolic acetyl-CoA from acetate. *YEF1*, encodes an NADH kinase that converts cytosolic NADH to NADPH (30). *GUT2* converts G3P to DHAP and participates in the G3P shuttle for transfer of electrons from cytosolic NADH to mitochondrial NADH (30). Conversely, mutations in *NDE1* (encoding an alternative enzyme for cytosol/mitochondrial NADH exchange and known to affect activity of Gut2 (86, 91-93)) had an apparent increase in lipid accumulation. In sum, our fitness data are consistent with the known importance of the precursors acetyl-CoA, G3P, and NADPH for TAG biosynthesis. However, the interactions of NADH transfer and glycerol metabolism in *R. toruloides* deserve more detailed study, as our results stand in contrast to observations in *Y. lipolytica* that *GUT2* mutants had increased lipid accumulation (94). Furthermore, the predominant source of NADPH to supply fatty acid synthesis remains unexplored in *R. toruloides* (see supplementary text for further discussion).

Finally, *RTO4_16381*, a distant homolog of *H. sapiens PLIN1* (perilipin), was also essential for high lipid accumulation, consistent with its homologs known roles in lipid body maintenance and regulation of hydrolysis (86, 93, 95). Our data are in accordance with previous observations that protein RTO4_16381 (previously named Lpd1) localized to lipid droplets in *R. toruloides* and that a GFP fusion construct localized to lipid droplets when heterologously expressed in *S. cerevisiae* (94, 96-99). RTO4_16381 is depicted as localized to the lipid droplet in Figure 6, along with eleven other lipid droplet-associated proteins with high confidence lipid accumulation phenotypes or significant fitness defects on fatty acids. The products of these genes were observed in proteomic analysis of *R. toruloides* lipid droplets by Zhu et al. (55), except for RTO4_11043 (similar to human *BSCL2*) and DGA1 which have been localized to the lipid droplet in many other species (100-106).

### Diverse morphological phenotypes for lipid accumulation mutants

To further characterize the phenotypes of our lipid accumulation mutants, we performed differential interference contrast (DIC) and fluorescence microscopy. The mutants showed a variety of phenotypes with respect to both cellular and lipid droplet morphology. Eight examples are highlighted in Figure 7. While wild type cells most commonly had two lipid droplets of similar size, several high lipid accumulation mutants had qualitatively more cells with 3 or more lipid droplets (e.g. *MET14*Δ, Figure 7)) or cells with a single dominant droplet (e.g. *RAC1*Δ, Figure 7). *RAC1*Δ also had qualitatively larger, more spherical cells. A KDELC-likeΔ mutant with increased lipid accumulation also showed a defect in cell separation likely reflective of combined defects in lipid accumulation, secretion, and cell wall/septum formation. All strains had a wide variation in lipid droplet size, consistent with high variance in BODIPY intensity measured by flow cytometry (Figure 4 – figure supplement 2A). Most low-lipid strains appeared morphologically similar to wild type with smaller lipid bodies (Figure 7 – figure supplement 1). However, a *BSCL2*-likeΔ (seipin) mutant showed an even larger variation in droplet size than wild type, consistent with observations in *S. cerevisiae* mutants for the homolog *SEI1/FLD1* (107-110) and likely reflective of a conserved function in lipid droplet formation and efficient delivery of lipid biosynthetic proteins to the growing lipid droplet (111, 112). Autophagy mutants (*ATG2*Δ) had the most uniformly small lipid droplets in elongated cells with enlarged vacuoles. Overall, the morphological phenotypes we observed in *R. toruloides* are similar to a number of previous microscopic screens for altered lipid accumulation in diverse eukaryotes (113).

**Figure 7.**
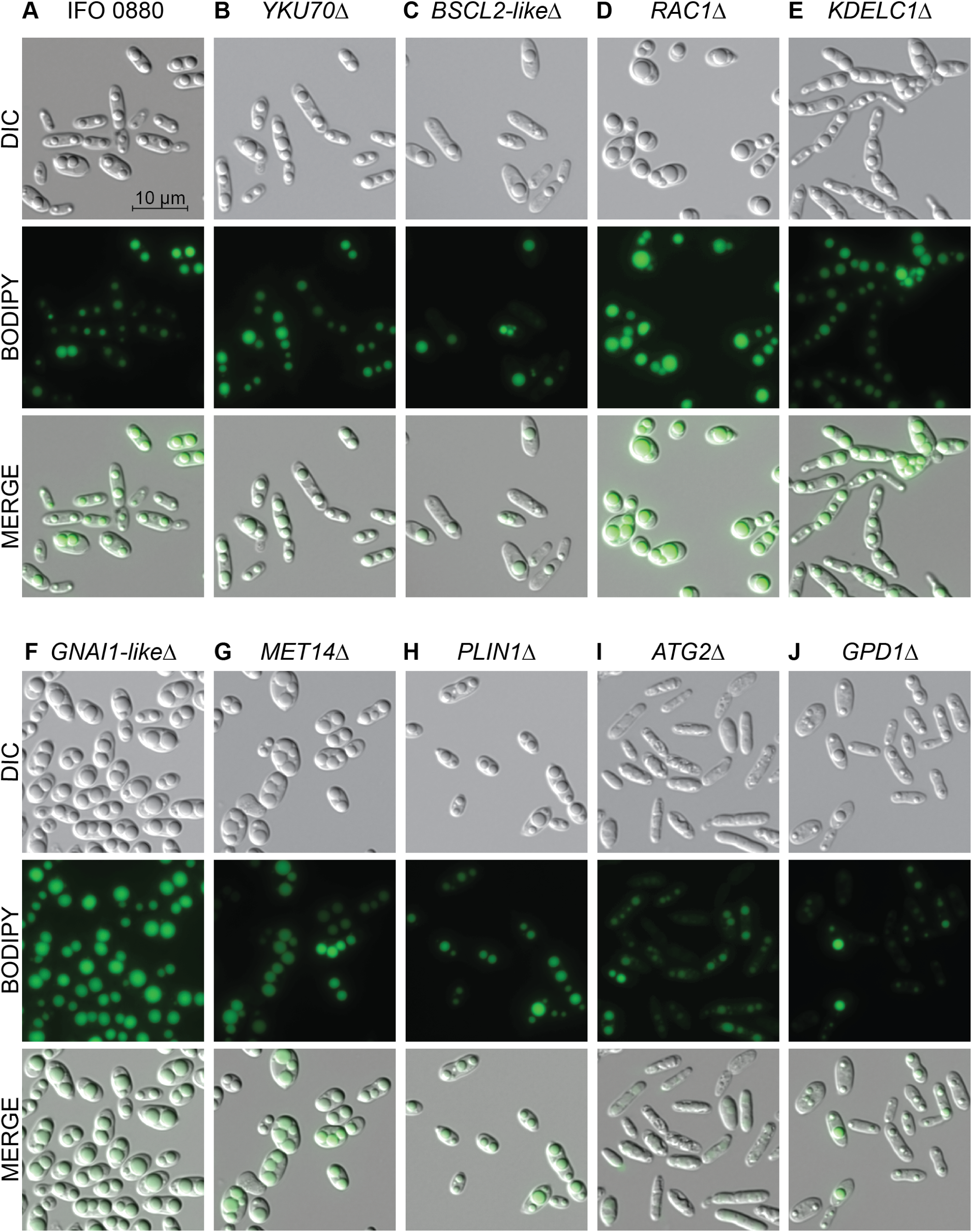
Light and fluorescence microscopy images of selected lipid accumulation mutants. DIC microscopy on eight deletion mutants for lipid accumulation genes. All deletion mutants (C-J) were constructed in a *YKU70*Δ background to enable homologous recombination at the targeted locus. Cells were grown 40 hours in low nitrogen lipid accumulation media. DIC, BODIPY 493/503 fluorescence, and composite images are shown for ten strains. (A) *R. toruloides* IFO 0880 (WT). (B) *RTO4J1920*Δ ortholog of *YKU70*. (C) *RTO4_11043*Δ similar to *H. sapiens BSCL2*. (D) *RTO4_14088*Δ ortholog of *H. sapiens RAC1*. (E) *RTO4_10371*Δ similar to *H. sapiens KDELC1*. (F) *RTO4_16215*Δ similar to *H. sapiens GNAI1*. (G) *RTO4_8709*Δ ortholog of *MET14*. (H) *RTO4_16381*Δ similar to *H. sapiens PLIN1*. (I) *RTO4_13598*Δ ortholog of *ATG2*. (J) *RTO4_12154*Δ ortholog of *GPD1*.

## Discussion

### Bringing functional genomics to non-model fungi with RB-TDNAseq

In this study, we employed a long-established method, *Agrobacterium tumefaciens*-mediated transformation, to extend barcoded insertion library techniques (8) into a non-model basidiomycetous fungi. We hope the wide range of species amenable to *A. tumefaciens* transformation (69, 114, 115) will allow RB-TDNAseq to be extended into fungal species for which it is not yet practical to construct random insertion libraries with other methods (e.g. transposon mutagenesis (116, 117) or *in vitro* transposition followed by homologous recombination (118)). We used RB-TDNAseq to map a sufficiently diverse set of insertion sites to measure the relative fitness of mutants in over 6,500 genes by tracking strain abundance in the mutant pool after competitive growth or physical enrichment using a simple, scalable BarSeq protocol. The fitness scores generated in our high-throughput experiments were consistent with predicted and measured fitness for single gene deletion strains. Also, because our genomic coverage is relatively complete, our insertion mapping also constitutes an initial survey of essential genes. Like all systematic surveys, this list is provisional, but we hope it will serve as a useful resource for genetics in *Rhodosporidium* species. These genes may also be potential targets for new antifungal strategies against basidiomycete pathogens, such as the closely related rusts of the Pucciniomycotina subphylum (119) and the more distantly related human pathogen *Cryptococcus neoformans* (117).

### New insights into fatty acid catabolism in *R. toruloides*

The presence of a probable mitochondrial fatty acid beta-oxidation pathway in *R. toruloides* has been noted previously (118). Our results confirm that this pathway is functional and essential for fatty acid utilization and add to mounting evidence that mitochondrial beta-oxidation is widespread in fungi (120). In mammals, some branched long-chain fatty acids are shortened in the peroxisome, then transferred via the acylcarnitine shuttle to the mitochondria for complete oxidation (44), while other long-chain fatty acids are metabolized solely in the mitochondria (121). *R. toruloides* has orthologs to the mammalian mitochondrial short, branched-chain and medium-chain acyl-CoA dehydrogenases *ACADSB* and *ACADM*, but not to the long-chain and very long-chain acyl-CoA dehydrogenases *ACADL* and *ACADVL*. Our observation that both peroxisomal and mitochondrial beta-oxidation were necessary for robust growth on fatty acids is consistent with conserved function for *ACADSB* and *ACADM* on short-chain fatty acids and a larger role for a diverse ensemble of peroxisomal acyl-CoA dehydrogenases and acyl-CoA oxidases in metabolism of longer-chain fatty acids. We also found that elements of the mitochondrial respiratory chain and amino acid biosynthesis were not essential for growth on glucose, but were necessary for robust growth on fatty acids. The importance of these pathways has been demonstrated in a gluconeogenic context in mammalian cells (122), so it remains unclear if their importance in *R. toruloides* amino acid catabolism is strictly in regards to efficient gluconeogenesis or if they also have more direct impacts on beta-oxidation.

We also observed differing fitness scores on oleic acid and ricinoleic acid for seven peroxisomal enzymes and three peroxins. We confirmed these phenotypes in deletion mutants for two acyl-CoA dehydrogenases (*RTO4_14567* and *RTO4_8963*) and the peroxin *RTO4_8673*. One hypothesis that would explain these differing phenotypes is divergent substrate specificity for the beta-oxidation enzymes, with different peroxins mediating effective localization of different enzymes. Both acyl-CoA dehydrogenases and acyl-CoA oxidases belong to an ancient superfamily that has been subject to a high rate of duplication, loss and lateral gene transfer suggesting a high rate of neofunctionalization (71-73). Different substrate specificity has been reported between orthologous mitochondrial acyl-CoA dehydrogenase in mice and humans (81, 123). Peroxisomal acyl-CoA oxidases showed overlapping, but distinct substrate specificity that even varied between different species of *Arabidopsis* (32). Our results demonstrate how a barcoded insertion library can accelerate discrimination of function between closely related members of a diversified gene family. These data may guide metabolic engineering strategies or the comprehensive biochemical assays necessary to characterize substrate specificity and enzymatic properties. Fitness assays on a much larger panel of substrates should yield further insights into the individual functions of *R. toruloides’* diverse complement of peroxisomal enzymes and guide experimental design for their biochemical characterization.

### Extending high-throughput fitness techniques to lipid production

While pooled fitness experiments have been used extensively to identify novel gene function, work so far has primarily focused on growth-based phenotypes, with only limited exploration of other phenotypes (8). In this study we used two proven strategies for differentiating between cells with altered lipid accumulation, buoyant density centrifugation (30) and FACS (92, 94, 96, 124-128), and applied them to our barcoded mutant pool. Inconsistencies between the two assays and with respect to independent BODIPY staining of targeted deletion strains suggests significant false positive rates for each assay in isolation. When both assays were in agreement, however, 17 of 21 deletion mutants had the expected phenotype in independent experiments. This approach identified 150 high confidence candidate genes with strong impacts on lipid accumulation under nitrogen limitation. While this set is likely incomplete, it complements previous transcriptional and proteomic studies to establish critical genes and cellular processes supporting lipid accumulation that deserve more intensive study. As has been noted in previous functional screens (99, 129-132), there was limited overlap between genes for which mutants had a detectable lipid accumulation phenotype in our study and genes with altered protein abundance in *R. toruloides* during lipid accumulation (35, 79, 97, 133, 134) (14 genes) or genes that co-purified with *R. toruloides* lipid droplets (5 genes) (98, 135, 136). The different ensemble of genes identified by each technique illustrate that these systems-level approaches complement each other, they do not replace each other.

### New insights into regulation of lipid metabolism in *R. toruloides*

Proteomic, transcriptomic, mutagenic and over-expression surveys of lipid metabolism have been carried out in several model eukaryotic systems including S. *cerevisiae* (91, 137-139), *C. elegans* (12-14), *D. melanogaster* (95, 96), various mammalian cell lines (31), and *Y. lipolytica* (140) (see Supplementary file 5 for a summary of genes identified in 35 studies). While wide variations in analytical techniques, nutrients, and culture conditions as well as a diverse genetic space make systematic comparisons between these surveys extremely difficult, a few broad themes are apparent. Protein trafficking and organelle interaction are inextricably linked with lipid body formation, growth and mobilization. Membrane-bound G proteins in the endomembrane network have conserved roles regulating trafficking and cellular morphology in response to metabolic states. A complex network of signaling cascades, protein modifications and transcription factors mediate the transition to lipid accumulation or lipid mobilization. A major output of this regulation is amino acid metabolism. Lipid metabolism and autophagy are deeply linked in a complex manner. Our findings were consistent with these general themes, including some orthologs to genes identified in the studies above, but the importance of general functions was more conserved across species than the roles of specific orthologous gene sets. The genes and processes we identify here should be considered in any strategy to optimize lipid metabolism in *R. toruloides* specifically or oleaginous yeasts in general. Comparative study of these processes across diverse species in standardized conditions will likely be required to uncover which aspects are fundamental to lipid droplet accumulation, maintenance and variation, and which processes are integrated by specific regulatory circuits in a given organism.

#### Organelle interactions and protein localization

Long regarded as essentially inert spheres of lipid, eukaryotic lipid droplets have of late come to be recognized as complex, dynamic, organelles with unique proteomic content and regulated interaction with other organelles (79). In animal cells, seipin *(H. sapiens* BSCL2) is thought to mediate lipid droplet nucleation from the ER (141). The *BSCL2* homolog *SEI1* was found to have conserved function in S. *cerevisiae* and *H. sapiens BSCL2* functionally complemented a *SEI1*Δ mutant. Cells with abnormally small and abnormally large lipid droplets were also reported in an *SEI1Δ* mutant (142). We found evidence that the closest *R. toruloides* homolog for *BSCL2* (*RTO4_11043*) has conserved function, as deletion mutants had quantitatively lower TAG content (as measured by flow cytometry) and qualitatively more cell-to-cell variation in lipid droplet sizes (by microscopy) than control strains. Perilipins (*H. sapiens* PLIN1-5) act as gatekeepers to the lipid droplet, regulating access by lipases (143) and possibly mediating interaction with mitochondria (144). Accordingly, we found that mutants for an *R. toruloides* perilipin homolog had reduced lipid accumulation. Protein trafficking between the ER and Golgi has been implicated in lipid droplet accumulation in *D. melanogaster*, specifically COPI retrograde transport is necessary to limit storage in lipid droplets (145). We found that disruption of *ERP1, ERP2, EMP24*, or *BST1* (implicated in ER to Golgi transport in COPII vesicles (14)) led to increased lipid accumulation. It is unclear at this time if COPI and COPII have different functions in lipid body formation across different eukaryotes, or if differing components of ER to Golgi trafficking are more critical when lipids are synthesized *de novo* from glucose or incorporated from exogenous fatty acids. (Beller et al. (30) cultured *D. melanoganster* cells on oleic acid to maximize lipid droplet size). Increased lipid accumulation in mutants with defective COPII trafficking might also be a function of impaired protein quality control (146). *H. sapiens DNAJC3* is implicated in regulation of the unfolded protein response by controlling elongation factor 2 phosphorylation (147). The *DNAJC3* ortholog *RTO4_14088* was a high confidence candidate for decreased accumulation as well. These data are consistent with a hypothesis that interaction between protein sorting, quality control and the unfolded protein response play a role in regulating lipid accumulation through modulation of protein translation. Alternatively, delivery of specific proteins to the lipid droplet via the vesicular trafficking system may be critical to lipid droplet growth and maintenance, or the effects of mutations in the endomembrane network on the lipid droplet may arise from redirection of carbon flux through membrane lipids.

#### G protein and kinase signaling cascades

We identified 28 genes with high-confidence roles in lipid metabolism that are homologous to genes implicated in G protein–coupled kinase signaling cascades, including RAC, Ras and Rab family G proteins. Rab GTPases are implicated in several aspects of vesicular traffic (148) and are also thought to mediate droplet fusion and interaction with endosomes (76). Several Rab family members have been identified in lipid droplets in *R. toruloides* (149), *S. cerevisiae, D. melanogaster*, and mammals, though their functional roles there remain unclear. 14-3-3 family proteins are known to affect several cellular processes (150) including protein trafficking (30) and modulate activity of both G proteins (134) and kinases (30). Rac and Ras G proteins have diverse roles in regulating the actin cytoskeleton, cell proliferation, cell cycle progression and polarity (128) and tend to localize to cell membranes, interacting with lipid kinases and transmembrane receptors (151-153). Likely both Rac1 and Ras1 interact directly with the lipid body, as Rac1 was detected in *R. toruloides* lipid droplets during nitrogen starvation (64, 154-156) and the Ras1 ortholog Ras85D was detected in *D. melanogaster* lipid droplets (157, 158). We were unable to quantify fitness scores for *RHO1*, but that G protein was also found associated with lipid droplets in *R. toruloides* (157, 158) and *S. cerevisiae* (159, 160). Undoubtedly these G proteins and downstream kinases function in a complex network of specific interactions, likely with considerable rearrangement of interactions from those observed in other species (64, 161, 162). Mapping these signaling networks in *R. toruloides* will require significant effort, but deep regulatory understanding will likely be required to truly optimize engineered pathways in any oleaginous yeast.

#### Autophagy and protein turnover

In mammalian and fungal cells, inhibition of autophagy has been reported to both decrease (64) and increase (161) lipid content. These discrepancies may be reflective of competing roles in fatty acid mobilization from lipid droplets and lipid droplet biogenesis, with different processes dominating in different cell types and under different conditions. Mechanisms of fatty acid mobilization have been proposed involving a macroautophagy-like process called lipophagy (17), a microautophagy-like process (microlipophagy) (8, 84), and autophagy-independent lipolysis (84). Why autophagy might be necessary for lipid droplet biogenesis is less clear, but autophagy-dependent recycling of membrane lipids to the lipid body has been demonstrated in mouse hepatocytes (162). Conversely, autophagy was also inhibited when TAG hydrolysis was impaired in HeLa cells (163) and when TAG synthesis or hydrolysis was blocked in S. *cerevisiae* (164) suggesting that these processes influence each other in a bi-directional manner. In both *Y. lipolytica* and *R. toruloides* several autophagy genes were transcriptionally induced under nitrogen starvation, co-incident with lipid accumulation (8). Further, in *Y. lipolytica*, chemical inhibition of autophagy strongly reduced lipid accumulation (165). In *S. cerevisiae* deletion of *ATG8* reduced lipid content, but that effect was lipolysis-dependent and *ATG3, ATG4, and ATG7* mutants were unchanged in lipid content (166).

Our findings demonstrated that autophagy was required for robust lipid accumulation in *R. toruloides*. While we cannot rule out a more direct role in lipid droplet growth and maintenance, a simple theory for this requirement is that autophagy is required for extensive recycling of cellular resources during lipid accumulation. Not only were several core components of autophagy necessary, but also the vacuolar proteases, and several proteins with predicted function in ubiquitination of proteins for proteosomal degradation. The methylcitrate cycle was required for robust lipid accumulation, which may be reflective of its proposed role in threonine recycling (167) or metabolism of propionyl-CoA from released odd-chained fatty acids (83, 168). How and why the role of autophagy in lipid droplet development varies by species and condition remains an open question, but *R. toruloides* is an attractive species in which to explore and answer those questions.

#### Amino acid biosynthesis and lipid accumulation

We also noted that disruption of several amino acid biosynthesis genes, particularly genes involved in sulfate assimilation into cysteine led to increased lipid production. These data are consistent with the repression of amino acid biosynthesis genes observed in *R. toruloides* (82) and other oleaginous fungi (167) in nutrient limited conditions. Notably, mutants for genes involved in methionine biosynthesis but not required for sulfate assimilation did not have enrichment scores reflective of increased lipid accumulation, nor did several arginine biosynthesis genes, or other auxotrophic mutants such as insertions in *PHA2* or *ADE5*. Mutants for *ARG1* had higher lipid content, but other mutants in the arginine pathway either had mixed results between the buoyancy and FACS assays *(ARG5* and *ARG7*), T-statistics below our thresholds *(ARG2, ARG7*, and *ARG8*) or showed no sign of increased lipid content *(CPA1, CPA2*, and *ARG3*). These discrepancies suggest that the increased lipid accumulation observed for some mutants may not be simply attributable to redirection of carbon flux from amino acid biosynthesis, but might be the result of active regulation in response to specific amino acids or metabolic intermediates. The transcriptional and proteomic response during nitrogen limitation in these mutants warrants deeper study.

#### tRNA thiolation, protein expression and carbon flux in nutrient limited conditions

Posttranslational modification of tRNAs has long been known to be critical to efficient protein translation in general (83), but in recent years thiolation of the U34 base on tRNAs for lysine (UUU), glycine (UUG), and glutamate (UUC) has been recognized to play an important role in fungal metabolic regulation generally (40) and particularly in response to stress such as nutrient limitation (169) and heat shock (27). In S. *cerevisiae*, defects in tRNA thiolation significantly alter protein expression for a large number of genes, but the mechanism of that change is disputed. Both transcriptional (170) and translational (53) mechanisms have been proposed. A commonality in these studies, however, is the altered expression of genes related to amino acid biosynthesis, protein expression and carbon metabolism. We found that any disruption in the URM1/elongator complex or tRNA thiolation process reduced lipid accumulation in our experimental conditions. The dramatic metabolic changes entailed in lipid accumulation under nutrient limitation may make for an informative framework in which to explore the mechanisms by which tRNA thiolation interacts with cellular metabolism.

#### Uncovering function for novel genes

In this study, we identified 46 *R. toruloides* genes with little or no functional predictions (Supplementary file 1), but which had important function in lipid metabolism as evidenced by reduced fitness when grown on fatty acids or altered lipid accumulation. These included 9 genes with broad conservation across ascomycete and basidiomycete fungi and 7 genes with conservation across several basidiomycete species. These genes are of particular interest for further study into their specific functions in lipid metabolism. Moreover, the mutant pool generated in this study should be an excellent tool to generate hypothetical functions for these genes and uncharacterized *R. toruloides* gene in general. Because the T-DNA insertions are barcoded, fitness experiments are inherently scalable to a large number of conditions. In this study, we examined a targeted set of conditions specifically aimed towards understanding lipid metabolism; however, we expect that generating a large compendium of fitness data in diverse conditions will enable a systematic survey of gene function in *R. toruloides* combining condition-specific data and cofitness analysis (53). We encourage the *R. toruloides* and broader fungal community to make use of this new resource.

#### Cell-to-cell variation in lipid accumulation

We noted extreme cell-to-cell variation in total lipid content in wild-type and mutant strains. This variation was evident in BODIPY fluorescence intensities that varied over at least an order of magnitude within any given sample (Figure 4 – figure supplement 2) and a wide range of lipid droplet sizes visible in microscopy images (Figure 7). Extreme variation in lipid accumulation is typical across eukaryotes, and has emerged as a useful paradigm to explore phenotypic diversity within isogenic populations (36). Our results indicate that *R. toruloides* may make a convenient system to dissect the genetic basis of single-cell phenotypic variation.

## Conclusions

In conclusion, we believe that RB-TDNAseq holds great promise for rapid exploration of gene function in diverse fungi. Because ATMT has been demonstrated in numerous, diverse fungi, we expect this method will be portable to many non-model species. Because the fitness analysis is inherently scalable, it will enable rapid fitness analysis over large compendia of conditions. Cofitness analysis of such compendia will accelerate the annotation of new genomes and identify new classes of genes not abundant in established model fungi. In this study, we demonstrated the application of RB-TDNAseq to the study of lipid metabolism in an oleaginous yeast that has significant potential to become a new model system for both applied and fundamental applications. We identified a large set of genes from a wide array of subcellular functions and compartments that impact lipid catabolism and accumulation. These processes and genes must be considered and addressed in any metabolic engineering strategy to optimize lipid metabolism in *R. toruloides* and other oleaginous yeasts. Deeper understanding of the extreme cell-to-cell variation in lipid accumulation seen across eukaryotes will likely require deeper mechanistic understanding of these processes and their interaction with the lipid droplet. The principles learned from exploring lipid metabolism and storage across diverse eukaryotes will inform biotechnological innovations for the production of biofuels and bioproducts, as well as new therapies for metabolic disorders.

## Acknowledgements

We thank Christopher Rao and Shuyan Zhang for initial advice and protocols for ATMT. We thank Kelly Wetmore and Adam Deutschbauer for their guidance on technical aspects of TnSeq and BarSeq experiments. We thank Morgan Price for his assistance and advice on TnSeq and BarSeq analysis, as well as for hosting our data on the fitness browser.

This material is based upon work supported by the U.S. Department of Energy, Office of Science, Office of Biological and Environmental Research program under Award Number DE-SC-0012527. Preliminary work establishing genetic, culturing, and assay protocols with *R. toruloides* was funded by grants OO1605 and OO6J01 from the Energy Biosciences Institute at the University of California Berkeley. Work performed at the DOE Joint BioEnergy Institute (http://www.jbei.org) is supported by the U.S. Department of Energy, Office of Science, Office of Biological and Environmental Research, through Contract No. DE-AC02-05CH11231 between Lawrence Berkeley National Laboratory and the U.S. Department of Energy. The work conducted by the U.S. Department of Energy Joint Genome Institute (http://jgi.doe.gov/), a DOE Office of Science User Facility, is supported by the Office of Science of the U.S. Department of Energy under Contract No. DE-AC02-05CH11231 between Lawrence Berkeley National Laboratory and the U.S. Department of Energy.

This work used the Vincent J. Coates Genomics Sequencing Laboratory at UC Berkeley, supported by NIH S10 Instrumentation Grants S10RR029668, S10RR027303, and S10OD018174.

## Methods

### Strains

We used *R. toruloides* IFO 0880 (also called NBRC 0880, obtained from Biological Resource Center, NITE (NBRC), Japan) as the starting strain for all subsequent manipulations. We used *Agrobacterium tumefaciens* EHA 105 and plasmids derived from pGI2 (36) for *A. tumefaciens* mediated transformation (ATMT) of *R. toruloides* (strain and plasmid kindly provided by Chris Rao, UIUC). The barcoded mutant pool was constructed by ATMT. We made all gene deletions in an non-homologous end-joining deficient *YKU70Δ* background (171) by homologous recombination of a nourseothricin resistance cassette introduced by either ATMT or electroporation of a PCR product. For deletions made by ATMT we used flanking arms of ~1000-1500 bp for homologous recombination. We found that as few as 40 bp of flanking sequence were sufficient for homologous recombination of PCR products at many loci. All strains used in this study, and primers used for strain construction and verification are listed in Supplementary file 4.

### Culture conditions

For most experiments, we used optical density (OD) as measured by absorbance at 600 nm on a Genesys 20 spectrophotometer (ThermoFisher Scientific 4001-000) as a metric for growth and to control inoculation density. For IFO 0880 grown in rich media, 1 OD unit represents approximately 30 million cells/mL. Unless otherwise noted, cultures were grown at 30 °C in 100 mL liquid media in 250 mL baffled flasks (Kimble 25630) with 250 rpm shaking on an Innova 2300 platform shaker (New Brunswick Scientific 39-M1191-0000) with constant illumination using a LumaPro 6W led lamp (Grainger 33L570). We used yeast-peptone-dextrose media (YPD, BD 242820) for general strain maintenance and rich media conditions. For auxotrophy experiments we used 0.67% w/v yeast nitrogen base (YNB) w/o amino acids (BD 291940) with 111 mM glucose (Sigma G7528) as our defined media and supplemented with 75 mM L-methionine (Sigma M9625), 75 mM L-arginine (Sigma A5006), or 0.2% w/v drop-out mix complete, which contains all 20 amino acids, adenine, uracil, p-Aminobenzoic acid, and inositol (US Biological D9515). To test growth and fitness on oleic acid (Sigma 01008 and 364525), ricinoleic acid (Sigma R7257), and methylricinoleic acid (Sigma R8750), we used this same defined media formulation with 1% fatty acid (by volume) instead of glucose. For lipid accumulation experiments, we pre-cultured strains for two generations in YPD (OD 0.2 to OD 0.8) then washed them twice and resuspended them at OD 0.1 in low nitrogen medium; 0.17% w/v yeast nitrogen base (YNB) w/o amino acids or ammonium sulfate (BD 233520), 166 mM D-glucose, 7 mM NH4Q (Fisher S25168A), 25 mM KH_2_PO_4_ (Fisher P285-3), and 25 mM Na_2_HPO_4_ (Sigma S0876). This is the C:N 120 formulation from Nicaud et al. (54). Unless otherwise specified, cultures were harvested for lipid quantification or fractionation after 40 hours of growth and lipid accumulation.

### Genome sequencing and *de novo* assembly

To generate an improved genome assembly for IFO 0880 we prepared genomic DNA for PacBio RS II sequencing (Pacific Biosciences). Genomic DNA was purified using a two-step protocol, first using glass bead lysis and phenol-chloroform extraction, as previously described (55), followed by a QIAGEN Genomic-tip 100/G method (QIAGEN, 10243). All QIAGEN buffers were obtained from a Genomic DNA Buffer Set (QIAGEN, 19060). Briefly, the dry genomic DNA pellet was first resuspended in G2 buffer supplemented with 200 μg/mL RNase A (QIAGEN 19101) and 13.5 mAU/ml Proteinase K (QIAGEN 19131), incubated at 50 °C for one hour, and then loaded on a Tip-100 column. After three washes with QC buffer and elution with QF buffer, the DNA was precipitated with isopropanol and removed by spooling using a glass Pasteur pipet. The genomic DNA was washed with 70% ethanol and after air-drying, resuspended in EB buffer (pH 7.5). DNA concentration was determined by fluorometry (Qubit, ThermoFisher Scientific) and submitted to University of Maryland Genomics Resource Center for library preparation and sequencing. A 10 kb insert size selected (BluePippin, Sage Science) SMRTbell library was prepared and sequenced on a PacBio RS II platform using P4C2 chemistry and 10 SMRT cells. *De novo* assembly of 610,663 polymerase reads (mean subread length of 5,193 bp) was performed using SMRT Analysis version 2.3.0.140936 (http://www.pacb.com/support/software-downloads/) and the RS_HGAP_Assembly.3 protocol (HGAP3) using default settings except for a genome size of 20,000,000 bp. The final assembly contained 30 polished contigs (mean coverage of 131-fold) with a total genome size of 20,810,536 bp. Paired-end Illumina data (17,817,326 PE100 reads, (172)) was used for error correction using Pilon version 1.13 (https://github.com/broadinstitute/pilon). As expected, the most common type of correction (569 in total) was insertion or deletion of a nucleotide in homopolymer regions. The final error corrected scaffolds were annotated by JGI and submitted to Genbank under the accession LCTV02000000. Raw sequence data (PacBio and Illumina) has been deposited in the NCBI SRA (SRP114401 and SRP058059, respectively).

### RNA sequencing and analysis

To harvest RNA for improved gene model prediction, we inoculated *R. toruloides* into 50 mL cultures in M9 Minimal Salts Solution (BD Difco 248510), 2 mM MgSO_4_ (Sigma M7506), 100 μM CaCl2 (Sigma C5670), and Yeast Trace Elements Solution (88 μg/mL nitrilotriacetic acid, 175 μg/mL MgSO_4_ 7H_2_O, 29 μg/mL MnSO_4_ H_2_O, 59 μg/mL NaCl, 4 μg/mL FeCl_2_, 6 μg/mL CoSO_4_, 6 μg/mL CaCl_2_ 2H_2_O, 6 μg/mL ZnSO_4_ 7H_2_O, 0.6 μg/mL CuSO_4_ 5H2O, 0.6 μg/mL KAl(SO_4_)_2_ 12H_2_O, 6 μg/mL H_3_BO_3_, 0.6 μg/mL Na_2_MoO_4_ H_2_O), pH 7.0, with 2% glucose (Sigma D9434) or 10 mM p-Coumaric acid (trans-4-hydroxycinnamic acid; Alfa Aesar A15167), and incubated overnight at 30 °C with 200 rpm shaking. We harvested cultures at mid-log phase, centrifuged at 3000 RCF for 10 minutes at room temperature, removed the supernatant and flash-froze the cell pellet in an ethanol/dry ice bath and stored at -80 °C. We lyophilized pellets overnight in a FreeZone-12 freeze dry system (LabConco 7754030) and extracted total RNA with a Maxwell RSC Plant RNA Kit (Promega AS1500) using a Maxwell RSC instrument (Promega AS4500). RNA was sequenced and mapped to the *R. toruloides* IFO 0880 genome at the Department of Energy Joint Genome Institute (JGI) in Walnut Creek, CA with in-house protocols.

### Gene model predictions and curation

The improved genome assembly was annotated using the JGI Annotation pipeline (38). Owing to relatively small intergenic spacing in the *R. toruloides* genome, fused gene models were a common problem. We hand curated over 500 gene models by searching for homology to unrelated proteins at each end of the automated gene models and inspecting agreement with assembled transcripts from our RNAseq experiments. Briefly, for all protein models over 400 amino acids long, we used the N-terminal and C-terminal 30% of each sequence in separate BLAST queries (NCBI BLAST-plus software 2.2.30) to a custom database of proteins from 22 other eukaryotic genomes (see Orthology relationships, below). We then compared the significant alignments for each terminus of a given gene and scored them for disagreement in regards to the respective orthology groups to which each target sequence belonged with a custom Python script (scripts and data files available at https://bitbucket.org/FungalTDNAseq/fusedgenemodels). The top-scoring 500 gene models were manually inspected for uncharacteristically long introns and for predicted introns and exons not supported by RNAseq reads and modified as required using the Mycocosm genome browser. The current genome annotation is publicly available at the JGI Mycocosm web portal (55): http://genome.jgi.doe.gov/Rhoto_IF00880_4

### Orthology relationships

We predicted orthologous proteins for our *R. toruloides* gene models in *H. sapiens, D. melanogaster, C. elegans, A. thaliana, C. reinhartii, S. cerevisiae*, and 16 other fungi with the orthomcl software suite version 2.0.9 (173). See Supplementary file 1 for a full list of ortholog groups and details on the genomes used in this analysis.

### Vector library construction

To efficiently construct a large and diverse mutant pool of barcoded mutants we first constructed a large library of barcoded vectors with an optimized type II-S endonuclease cloning strategy (38). We modified the ATMT vector pGI2 (Ref: Appl Microbiol Biotechnol (2013) 97:283-295, DOI 10.1007/s00253-012-4561-7) to act as a barcode receiving vector by first removing the two pGI2 SapI sites already present on the vector backbone through SapI restriction digestion, treatment with T4 DNA polymerase for blunt end formation and subsequent blunt end ligation. Next, we introduced two divergent SapI recognition sites just inside the right border of the T-DNA (vector pDP11) as the integration site for random barcoding. We added the barcodes by synthesizing the oligonucleotide GATGTCCACGAGGTCTCTNNNNNNNNNNNNNNNNNNNNCGTACGCTGCAGGTCGAC and amplifying with primers TCACACAAGTTTGTACAAAAAAGCAGGCTGGAGCTCGGCTCTTCGCCCGATGTCCACGAGGTCTCT and CTCAACCACTTTGTACAAGAAAGCTGGGTGGATCCGCTCTTCAATTGTCGACCTGCAGCGTACG. We then combined 4 μg of vector and 140 ng of barcode fragments in a 50 μl reaction with 5 μl 10x T4 ligase buffer, 5 μl 10x NEB Cutsmart buffer (NEB, B7204S), 2.5 μl T7 ligase (NEB, M0318L), and 2.5 μl of SapI (NEB, R0569S). We incubated the reaction at 37 °C for 5 minutes, then 25 cycles of 37 °C for 2 minutes and 20 °C for 5 minutes, before denaturing the enzymes for 10 minutes at 65 °C. Without cooling the product, we added 1 μl SapI and incubated for 30 minutes at 37C to digest any uncut vector, then cooled to 10 °C. We purified the barcoded plasmids using a Zymo DNA clean and concentrator kit (Zymo Research D4014), eluting in 15 μl of elution buffer and pooled 10 barcoding reactions. We then transformed *E. coli* electrocompetent 10 Beta cells (NEB) according to the manufacturers specifications in 30 independent transformations. We estimated the diversity of the barcoded vector pool by performing barcode sequencing as described below, sequencing on an Illumina MiSeq system and estimating the true pool size by the relative proportion of barcodes with 1 or 2 counts. See the script Multicodes.pl from Wetmore et al. (174) for details. This yielded a barcoded pool estimated to consist of ~100 million clones.

To efficiently construct a large and diverse mutant pool of barcoded mutants we first constructed a large library of barcoded vectors with an optimized type II-S endonuclease cloning strategy (38). We modified the ATMT vector pGI2 (Ref: Appl Microbiol Biotechnol (2013) 97:283-295, DOI 10.1007/s00253-012-4561-7) to act as a barcode receiving vector by first removing the two pGI2 SapI sites already present on the vector backbone through SapI restriction digestion, treatment with T4 DNA polymerase for blunt end formation and subsequent blunt end ligation. Next, we introduced two divergent SapI recognition sites just inside the right border of the T-DNA (vector pDP11) as the integration site for random barcoding. We added the barcodes by synthesizing the oligonucleotide GATGTCCACGAGGTCTCTNNNNNNNNNNNNNNNNNNNNCGTACGCTGCAGGTCGAC and amplifying with primers TCACACAAGTTTGTACAAAAAAGCAGGCTGGAGCTCGGCTCTTCGCCCGATGTCCACGAGGTCTCT and CTCAACCACTTTGTACAAGAAAGCTGGGTGGATCCGCTCTTCAATTGTCGACCTGCAGCGTACG. We then combined 4 μg of vector and 140 ng of barcode fragments in a 50 μl reaction with 5 μl 10x T4 ligase buffer, 5 μl 10x NEB Cutsmart buffer (NEB, B7204S), 2.5 μl T7 ligase (NEB, M0318L), and 2.5 μl of SapI (NEB, R0569S). We incubated the reaction at 37 °C for 5 minutes, then 25 cycles of 37 °C for 2 minutes and 20 °C for 5 minutes, before denaturing the enzymes for 10 minutes at 65 °C. Without cooling the product, we added 1 μl SapI and incubated for 30 minutes at 37C to digest any uncut vector, then cooled to 10 °C. We purified the barcoded plasmids using a Zymo DNA clean and concentrator kit (Zymo Research D4014), eluting in 15 μl of elution buffer and pooled 10 barcoding reactions. We then transformed *E. coli* electrocompetent 10 Beta cells (NEB) according to the manufacturers specifications in 30 independent transformations. We estimated the diversity of the barcoded vector pool by performing barcode sequencing as described below, sequencing on an Illumina MiSeq system and estimating the true pool size by the relative proportion of barcodes with 1 or 2 counts. See the script Multicodes.pl from Wetmore et al. (174) for details. This yielded a barcoded pool estimated to consist of ~100 million clones.

### *Agrobacterium* mediated transformation of *R. toruloides*

We transformed the barcoded vector pool into *A. tumefaciens* EHA 105 with a protocol adapted from established methods (174). We diluted a stationary phase starter culture 1:100 in 500 ml Luria-Bertani broth (BD 244620) and cultured for 6 hours at 30 °C. We pelleted cells at 3000 RCF, 10 minutes, 4 °C, washed pellets in ice-cold 1mM HEPES (Fisher Scientific, BP310), pH 7.0, then washed them in ice-cold 10% glycerol 1 mM HEPES, suspended cells in 5 ml ice-cold 10% glycerol 1mM HEPES, and flash froze 50 μl aliquots in liquid nitrogen. To produce a large transformant pool of *A. tumefaciens* bearing millions of unique barcode sequences, we electroporated 5 ml of competent cells with 50 μg of plasmid DNA in a BTX HT100 96-well plate chamber (50 μl per well) with a 2.5 kV pulse, 400 ohm resistance and 25 μF capacitance from a BTX ECM 630 wave generator. We recovered cells in LB for 2 hours at 30 °C, and plated on LB agar with 50 μg/ml kanamycin (Sigma, K4000). Approximately 14 million transformation events were scraped and collected into a mixed pool for transformation of *R. toruloides*.

We grew the barcoded *A. tumefaciens* pool to OD 1 in 50 mL YPD in a baffled flask at 30 °C, then pelleted the cells and suspended in 10 mL induction medium (1 g/L NH_4_CL, 300 mg/L MgSO_4_ 7H_2_0, 150 mg/L KCl (Fisher P267-500), 10 mg/L CaCl_2_ (VWR 0556), 750 μg/L FeSO_4_ 7H_2_O (Acros 423731000), 48 mg/L K2HPO4 (VWR 0705), 3.9 g/L NaH_2_PO_4_ (Fisher BP329), 198 mg/L D-Glucose, 1 mg/L thiamine (Sigma T4625), and 196 μg/L acetosyringone (Sigma, D134406)) and incubated 24 hours at room temperature in culture tubes on a roller drum. We cultured *R. toruloides* in 10 mL YPD to OD 0.8, then pelleted the cells and suspended in the induced *A. tumefaciens* culture for 5 minutes at room temperature. We filtered the mixed culture on a sterile 0.45 μm membrane filter (Millipore, HAWP04700) then transferred the filter to induction media 2% agar (BD 214010) plates for incubation at 26 °C for 4 days. We then washed the filters in sterile water and plated on YPD 2% agar with 300 μg/ml cefotaxime (Sigma C7039) and 300 μg/ml carbenicillin (Sigma C1389) and incubated at 30 °C for two days. We scraped these plates to collect transformed *R. toruloides*, recovered the mutant pool in YPD plus cefotaxime and carbenicillin for 24 hours, added glycerol to 15% by volume and stored at -80 °C. We repeated this protocol 40 times to recover approximately 2 million transformation events. In some rounds of transformation, we also included 0.05% casamino acids (BD 223120) or 1% CD lipid concentrate (Gibco 11905-031) in the induction media plates to promote recovery of mutants with impaired amino acid or lipid biosynthesis. We then recovered each of these transformation subpools on YPD plus cefotaxime and carbenicillin 12 hours to clear residual *A. tumefaciens* and combined them into one master pool, divided it into 1 ml aliquots in YPD 15% glycerol and stored them at -80 °C.

### TnSeq library preparation

To isolate high quality genomic DNA we harvested ~10^8^ cells from a fresh YPD culture of the mutant pool, washed the pellet in water and suspended in 200 μl TSENT buffer (2% Triton X-100 (T8787-50ML), 1% SDS (Ambion AM9820), 1 mM EDTA (Sigma ED2SS), 100 mM NaCl (Sigma S5150), 10 mM Tris-HCl, pH 8.0 (Invitrogen 15568-025)). We then added the sample to 200 μl 25:24:1 phenol/chloroform/isoamyl alcohol (Invitrogen 15593-031) in screw-top tubes with glass beads (Benchmark 1031-05) on ice and vortexed for 10 minutes at 4 °C. We added 200 μl TE buffer (Ambion AM9858), centrifuged 20 minutes at 21,000 RCF 4 °C, removed the aqueous phase to 1 mL ethanol (Koptec V1016) and centrifuged 20 minutes at 21,000 RCF at 4 °C to pellet DNA. DNA was dried and suspended in 200 μl TE, treated with 0.5 μl RNase A (Qiagen 19101), then purified with the Zymo Research Genomic DNA Clean and Concentrator Kit (D4064). We checked DNA quality on a 0.8% agarose E-Gel (Thermo Scientific G51808) and quantified with a Qubit 3.0 fluorometer using the dsDNA HS reagent (Invitrogen 1799096).

To sequence sites of genomic insertions we followed the TnSeq protocol of Wetmore et al. (55), using their Nspacer_barseq_universal primer and P7_MOD_TS_index primers for final amplification (Supplementary file 4). Because we found a high proportion of non-specific products in our TnSeq mapping and highly variable recovery of the same insertions between technical replicates, we sequenced multiple replicates for each batch of ATMT mutants (around 10,000 – 100,000 mutants per batch) and used at least two annealing temperatures for the final PCR enrichment for each batch. In total, we sequenced about 900 million reads from 64 independent TnSeq libraries. A full summary of TnSeq libraries used to map the mutant pool is listed in Supplementary file 4. Libraries were submitted for single-end 150 bp Illumina sequencing on a HiSeq 2500 platform at the UC Berkeley Vincent J. Coates Genomics Sequencing Laboratory, except for subset of smaller runs on an Illumina MiSeq platform as indicated in Supplementary file 4. Sequence data have been submitted to the NCBI Short Read Archive (SRP116146).

### Mapping insertion locations

We used a similar strategy as Wetmore et al. (39) to map the location of each barcoded T-DNA insertion, with minor alterations. Our modified code is available at: (https://bitbucket.org/FungalTDNAseq/rb-tdnaseq)

MapTnSeq_trimmed.pl processes the TnSeq reads to identify the barcode sequence and is a modified version of MapTnSeq.pl (62), with three minor alterations. We ignore the last 10 bases of the T-DNA sequence, as the length of T-DNA border sequence included in the final insertion is variable. We allow for barcode sequences of 17 – 23 basepairs instead of exactly 20. We report all TnSeq reads in which sequence past the end of the expected T-DNA insert aligns with other regions of the T-DNA sequence, or with the outside vector as ‘past end’ reads. These are mappings of junctions between concatemeric T-DNA inserts and unprocessed T-DNA vectors, respectively.

RandomPoolConcatemers.py is a custom script that associates barcode sequences mapped in MapTnSeq_trimmed.pl with genomic locations and then filters those barcodes for insertions at unique, unambiguous locations. First, for all barcodes sequenced, the number of reads mapping to any genomic location and the number of reads mapping to concatemeric junctions are tabulated. Any barcodes that only differ by a single basepair from a barcode with 100 times more reads are removed as likely sequencing errors and reported as ‘off by one’ barcodes. Any barcode for which there are more than 7 times as many ‘past end’ reads as reads mapping to genomic locations as ‘past-end’ barcodes. The past-end barcodes are further characterized as ‘head-to-tail’ concatemers (majority of Tnseq reads map to the left border T-DNA sequence), ‘head-to-head’ concatemers (majority of the reads map to the right border T-DNA sequence), or ‘Runon’ insertions (majority of reads map to pGI2 outside the T-DNA sequence). Any barcodes for which the majority of TnSeq reads map ambiguously to the genome are removed and reported as ambiguous barcodes. Any barcodes for which 20% or more of the TnSeq reads map to a different location than the most commonly observed location are removed and reported as ‘multilocus’ barcodes. Finally, any barcodes mapped within 10 bases of a more abundant barcode for which there is a Levenshtein edit distance (55) less than 5 are removed as likely sequencing errors and reported as ‘off by two’ barcodes. The remaining unfiltered barcodes are reported as the mutant pool.

InsertionLocationJGI.py is a custom script to match the genomic locations of barcodes in the mutant pool to the nearest gene in the current JGI *R. toruloides* gene catalog and report whether the insertion is in a 5-prime intergenic region, a 5-prime UTR, an exon, an intron, a 3-prime UTR, or a 3-prime intergenic region of that gene.

InsertBias.py is a custom script to analyze potential biases in T-DNA insertion rates. The script tracks number of insertions versus scaffold length for all scaffolds in the genome, GC content in the local regions of insertion, and insertion rates in promoter regions, 5-prime untranslated mRNA, exons, introns, 3-prime untranslated mRNA, and terminator regions. To assess fine-scale biases in insertion locations, all locations in the genome are apportioned to one of the above feature types, then for each feature type, the same number of insertions as were observed for that feature type in the mutant pool are sampled at random (without replacement) from all the genomic locations assigned to that feature type.

### Barcode sequencing

We isolated genomic DNA with the Zymo Research Fungal/Bacterial DNA MiniPrep kit (D6005). We used Q5 high-fidelity polymerase with GC-enhancer (New England Biolabs M0491S) to amplify unique barcode sequences flanked by specific priming sites, yielding a 185 bp Illumina-sequencing-ready product (Figure 1 – figure supplement 1). We used BarSeq primers from Wetmore et al. (175) (Supplementary file 4), except we replaced primer P1 with a mix of primers with 4-6 random bases to improve nucleotide balance for optimal sequencing of low-diversity sequences (174). We cleaned PCR products with the Zymo Research DNA clean and concentrator kit (D4014). We quantified product yield with a Qubit 3.0 fluorometer system and mixed as appropriate for sequencing as multiplexed libraries. We sequenced libraries on an Illumina HiSeq 4000 system at the UC Berkeley Vincent J. Coates Genomics Sequencing Laboratory. Libraries were purified with a Pippin Prep system (Sage Biosciences) and loaded with 20% PhiX DNA as a phasing control for low diversity samples (174). We sequenced each biological replicate to a depth of at least 20 million reads. We counted occurrences of T-DNA barcodes in each sample with the script MultiCodes_Variable_Length.pl, a modified version of MultiCodes.pl from Wetmore et al. (38) that allows for barcodes of 17 – 23 basepairs.

### Fitness analysis

For all BarSeq experiments, we thawed frozen aliquots of the mutant pool on ice and inoculated them into YPD at OD 0.2. Cultures were recovered approximately 12 hours until OD 600 was approximately 0.8. Cultures were pelleted at 3000 RCF for 5 min, washed twice in the appropriate media, and transferred to the condition of interest. Samples were taken from the YPD starter cultures (T_0_) and after 5-7 doublings in the experimental condition (T_condition_). Average fitness scores and T-like statistics (T-stats) as metrics for consistency between individual insertion mutants in each gene were calculated with Wetmore et al’s software (176) (combineBarSeq.pl and FEBA.R). Because that software does not consider biological replication between independent cultures, we then averaged fitness scores for each condition and combined T-stats across replicates with the script AverageReplicates.py, treating them as true T-statistics. That is: T_condition_ = Sum(T_replicates_)/Sqrt(N_replicates_). T-stats computed in this way give a measure of significance for observed fitness for each gene with respect the total population in that condition. To assess significance of differences in observed fitness between growth conditions we computed T_c1 – c2_ = (F_c1_ – F_c2_) / Sqrt ((F_c1_/T_c1_)^2^ + ((F_c2_/T_c2_)^2^) with the script ResultsSummary.py. For experiments performed simultaneously with explicitly paired samples (e.g. biological replicates in two or more conditions that originated from the same T_0_ sample), we also computed an alternative statistical test, the Wilcoxon signed rank test (177, 178) with custom software (Wilcoxon.py). Briefly, this program takes the raw counts for each barcode in an experiment, normalizes them by sequencing depth per sample, then groups barcodes disrupting the same gene and performs the Wilcoxon signed rank test on the difference between normalized counts for all barcodes in the paired samples. For consistency with Wetmore et al.’s (179) algorithms, we only included data from barcodes disrupting the central 80% of the coding region and with at least 3 counts in one condition in the signed rank tests. We generated K-means clusters of fitness scores using Pearson correlation as the similarity metric using Cluster 3.0 (53). For comparing enrichment in density and FACS separated fractions we computed F and T for each fraction versus the T_0_ control. The enrichment score E and T between fractions was then calculated as E = F_high lipid_ – F_low lipid_ and T_high lipid – low lipid_ = (F_high lipid_ – F_low lipid_)/ Sqrt ((F_high lipid_/T_high lipid_) + ((F_low lipid_/T_low lipid_)) with the script ResultsSummary.py. We generated hierarchical clusters of enrichment scores using Pearson correlation as the similarity metric and average linkage as the clustering method. All fitness data are available in Supplementary file 2 and the fitness browser ((180)). Custom Python scripts are available at ((180)). Sequence data have been submitted to the NCBI Short Read Archive (SRP116193)

### Transformation of *R. toruloides* by electroporation

We cultured *R. toruloides* overnight in 10 mL YPD on a roller drum to an OD 600 of 2, then pelleted cells at 3000 RCF, 5 min at 4 °C in a benchtop centrifuge (Eppendorf 5810 R). Cells were kept at 4 °C from this point. We transferred the pellets to 1.5 mL tubes and washed them 4 times with ice cold 0.75 M D-sorbitol (Sigma S1876), centrifuging each wash 30 seconds at 8000 RCF, 4 °C (Eppendorf 5424). After the final wash, we removed excess D-sorbitol and added 35 μl of cell pellet to 10 μl of fresh 0.75 M D-sorbitol and ~1 μg of PCR product in 5 μl water in a chilled 0.1 cm cuvette. We electroporated cells at 1500 kV, 200 ohms and 25 μF with a ECM 630 (BTX) electroporation system. We then added 1 mL cold 1:1 mixture of YPD and 0.75 M D-sorbitol and transferred to 14 mL round bottom culture tubes for a 3-hour recovery culture at 30C with shaking at 200 rpm on a platform shaker. We then pelleted the cultures at 8000 RCF, 30 seconds, suspended in 200 μl YPD and plated on YPD with 100 μg/mL nourseothricin (clonNAT, Werner Bioagents).

### Gene ontology enrichment

We scored enrichment of gene ontology terms with a custom script that performs a hypergeometric test on the frequency of each term in the genome versus the frequency in given gene set (script GOenrich.py, available at (181)). We corrected for multiple hypothesis testing with the Benjamini-Hochberg correction (176). We extended the GO terms associated with *R. toruloides* genes in the current JGI annotation by collecting terms for orthologous genes in *Arabidopsis thaliana, Aspergillus nidulans, Caenorhabditis elegans, Candida albicans, Homo sapiens, Mus musculus*, and *Saccharomyces cerevisiae*, obtained from the Gene Ontology Consortium (177, 178).

### Total fatty acid quantification with gas chromatography

Cell lysis, extraction of total lipids, and conversion to fatty acid methyl esters (FAMEs) was based on a published protocol (179). We cultured IFO 0880, a selection of seven targeted deletion strains (see Supplementary file 6) and one overexpression strain (RT880-AD, (53)) in low nitrogen medium for 48 or 96 hours. We collected paired 5 mL samples from each in screw-top glass tubes (Corning 99502-10) and 15 mL polyethylene tubes (Corning 352096) for lipid extraction and mass determination, respectively. We pelleted samples by centrifugation at 2000 RCF, 4 °C for 20 minutes, and washed once in water to remove salts and unused glucose. We then transferred the mass determination sample to a pre-tared 1.5 mL microcentrifuge tube. We froze both samples at -20 °C overnight, then lyophilized them 48 hours in a FreeZone freeze dry system (Labconco 7754042) before weighing/extraction. We added 1 mL methanol spiked with 250 μg methyl tridecanoate to each sample to serve as an internal standard (ISTD). We then resuspended lipid extraction samples (usually about 10-20 mg) by vortexing in 3 mL 3N methanolic HCL (SUPELCO 33050) and 200 μl chloroform (Sigma 472476) and incubated at 80 °C water bath for 1 hour. Cell lysis and conversion to FAMEs occurs during this incubation. To extract FAMEs we then added 2 mL hexane (Sigma 650552) and vortexed samples well before centrifugation at 3000 RCF for 3 minutes. One μL of the hexane layer was injected in split mode (1:10) onto a SP2330 capillary column (30 m x 0.25 mm x 0.2 μm, Supelco). An Agilent 7890A gas chromatograph equipped with a flame ionization detector (FID) was used for analysis with the following settings: Injector temperature 250 °C, carrier gas: helium at 1 mL/min, temperature program: 140 °C, 3 min isocratic, 10 °C/min to 220 °C, 40 °C/min to 240 °C, 5 min isocratic. FAME concentrations were calculated by comparing the peak areas in the samples to the peak areas of ten commercially available high-purity standards (C16:0, C16:1, C17:0, C18:0, C18:1, C18:2, C20:0, C20:1, C22:0, C24:0) (Sigma) in known concentration relative to the internal standard, respectively.

### Relative TAG measurement with BODIPY and flow cytometry

We inoculated deletion mutants and the *YKU70*Δ parental strain at OD 0.1 in low nitrogen medium and cultured for 40 hours. We fixed samples by adding 180 μl cell culture to 20 μl 37% formaldehyde (Electron Microscopy Sciences) and incubating for 15 minutes at room temperature. We then diluted fixed cells 1:100 in 200 μl PBS (from 10X concentrate, Gibco 70011-44) with 0.5 M KI and 0.25 μg/mL BODIPY 493/503 (Life Technologies D-3922), then incubated 30 minutes at room temperature. We quantified BODIPY signal for 10,000 cells per sample on a Guava HT easyCyte system (EMD Millipore) in the green channel (excitation 488 nm, emission 525 nm) using InCyte software (Millipore).

### Population enrichment with FACS

We cultured the barcoded mutant pool in low nitrogen medium for 40 hours. We then diluted unfixed cells 1:100 in 10 ml PBS with 0.5 M KI and 0.25 μg/mL BODIPY 493/503, then incubated 30 minutes at 30 °C with shaking. We then sorted the population on a Sony SH800 cell sorter with a 70 μM fluidic chip, sorting in semi-purity mode. We first applied a gate for single cell events with forward scatter height within 15% of forward scatter area. We sorted a sample of 10 million cells with the scattering gate alone as a control population, to account for effects of growth, sorting, and collection that are independent of lipid accumulation. Then we collected the 10% of the size-filtered population with the highest and lowest signals in the FITC channel. We collected 10 million cells each for the high and low signal populations. We collected all sorted cells in YPD with 300 μg/ml cefotaxime (Sigma C7039) and 300 μg/ml carbenicillin (Sigma C1389), then grew them to saturation in our standard culture conditions and pelleted 1 mL sample, and then stored at -20 °C for BarSeq analysis.

### Population enrichment with sucrose density gradients

We prepared linear sucrose gradients with the method of Luthe et al (180). For example, to prepare a 65%-35% sucrose gradient; we prepared four solutions of sucrose (Sigma G7528) at 65, 55, 35, and 35 grams per 100 mL in PBS, then successively froze 10 mL layers of each concentration in a 50 mL conical tube (Corning 430829) on dry ice and stored the gradient at -20 °C. We selected appropriate gradients to maximize the physical separation of the cell population by running trial experiments with wild type IFO 0880 cultures on a number of sucrose gradients. The gradients used in each experiment are described in Figure 4 – figure supplement 2. Approximately 24 hours before performing density separation on cell population, the appropriate step gradient was moved to 4 °C to thaw, yielding a linear gradient (180).

To perform the separation, we centrifuged 50 mL of culture at 6,000 RCF at 4 °C for 20 min. We then suspended the pellet in 5 ml PBS at 4 °C and carefully loaded it onto a sucrose gradient. We centrifuged the gradients for 1 hour at 5,000 RCF at 4 °C with slow acceleration and no brake for deceleration in a Beckman Coulter Avanti J26 XP centrifuge with a JS5.3 swinging bucket rotor. To collect fractions, we pierced the bottom of each tube with the tip of needle (BD PrecisionGlide 16G, 305197), to slowly drain the gradient from the bottom, at 1 drop every 1-5 seconds. We collected 2 mL fractions, estimated average fraction density by weighing a 100 μl sample (Figure 4 – figure supplement 2) and measured the distribution of the cell population across the sample by optical density (Figure 4 – figure supplement 1D). The appropriate fractions were then combined to sample the least buoyant (highest density) 5-10%, median buoyancy 30-50%, and most buoyant (lowest density) 5-10% of the population. For each biological replicate, we also collected a 1 mL sample from culture before separation to monitor growth in the experimental condition.

### Microscopy

Cover slips were submerged in 0.1% v/v polylysine (Sigma P8920) for 15 minutes. Cover slips were removed from polylysine and blotted dry from the bottom of vertically-held slips. Slips were then washed several times with ddH_2_O and rapidly dried with compressed air. Directly prior to imaging, slips are visually inspected for streaks and dust and softly cleaned with lens paper. Cells were grown 40 hours in low nitrogen medium 1 mL of culture was transferred to 2 mL microcentrifuge tubes with 1 mL of PBS and tubes were mixed briefly by vortexing. Cells were pelleted at 9000 RCF for 1 minute in a microcentrifuge, aspirated and suspended in 100 μl of fluorescent staining solution (PBS with 0.5 M KI and 0.25 μg/mL BODIPY 493/503) to visualize intracellular lipid droplets. Four μl of stained cells were pipetted up and down and transferred to the clean slides. Polylysine-coated cover slips were carefully placed on 4 μl drop to ensure even spreading of liquid. Cells were observed on an Axio Observer microscope (Zeiss) with a plan-apochromat 100x DIC objective (Zeiss 440782-9902-000), ORCA-Flash 4.0 camera (Hamamatsu C11440-22CU), and ZenPro 2012 (blue edition) software. For BODIPY imaging cells were illuminated with an X-cite Series 120 arc-lamp (EXFO Photonics Solutions) and 38HE filter set, 450-490 excitation, 500-550 emission (Zeiss 489038-9901-000). Zvi files were converted to 16 bit TIFF images and representative fields of view were cropped and channels merged using FIJI image processing software (181).

## Supplementary Text

### Refining the *R. toruloides* IFO 0880 genome sequence and annotation

An effective functional genomics approach requires high quality genomic sequence and reliable gene models. To improve assembly, we added long-read sequencing from Pacific Biosciences to our previously published data from Illumina sequencing (1). The refined gapless assembly is high quality, consisting of 21 megabases on 30 scaffolds (N50 = 6, L50 = 1.4 Mb) and a complete 112 Kb mitochondrial genome. Seven *de novo* scaffolds have telomeric repeats (2) at both ends, suggesting they represent complete chromosomes, and seven scaffolds have a telomeric repeat at one end (Supplementary file 1). For comparison, electrophoretic karyotyping of *R. toruloides* NP11 indicated 16 total chromosomes (3). We also used 100bp paired-end Illumina sequencing of mRNA to improve gene model prediction. The revised genome *(Rhodosporidium toruloides* IFO0880 v4.0) encoding 8490 predicted proteins is available at the Joint Genome Institute’s Mycocosm genome portal (4) and Genbank accession LCTV02000000. While the bulk of the gene models were predicted with the JGI’s automated protocols, erroneous fusion of neighboring genes was a significant issue. We have manually corrected several hundred fused models not supported by RNAseq data and encourage the *R. toruloides* research community to continue annotation refinement through the JGI portal. Summary tables of gene IDs, predicted functions, and probable orthologs in other systems are included in Supplementary file 1.

### Additional detail on mapping insertion locations with RB-TDNAseq

We adapted a high-throughput phenotyping strategy previously demonstrated in bacteria (5) by employing *Agrobacterium tumefaciens* mediated transformation (ATMT)(Figure 1A). Briefly, we created a large barcoded mutant pool in which *A. tumefaciens* transfer DNAs (T-DNA) bearing an antibiotic resistance cassette and a 20 base-pair random sequence (barcode) were inserted randomly throughout the genome. We then mapped the location of each insertion and its associated barcode with RB-TDNAseq, a variant of RB-TnSeq (a high-throughput method to enrich and sequence a diverse pool of transposon/genome junctions (5)), applied to T-DNA inserts. A more detailed view of the junction sequence and primers used for RB-TDNAseq and BarSeq are shown in Figure 1 – figure supplement 1.

From a mutant pool of approximately two million *R. toruloides* colonies, we sequenced 1,391,040 unique barcoded insertions with RB-TDNAseq. We successfully mapped 293,613 barcodes (21%) to T-DNA insertions at unique, unambiguous locations in the *R. toruloides* genome. The remainder of sequenced barcodes could not be mapped for several reasons (Figure 1 – figure supplement 2A). T-DNA is often inserted in concatemeric repeats (6-8), in which case only RB-TDNAseq reads from the terminal repeat provides mapping information. If the terminal repeat is truncated (9), or if it abuts genomic sequence that is recalcitrant to sequencing for any reason, then we are able to detect the barcode at junctions between T-DNA repeats, but not at junctions with the genome. 47% of sequence barcodes were not mappable for this reason. Likewise, if the terminal T-DNA is inserted in an inverted orientation, the result is an unmappable convergent concatemer (5% of barcodes). About 16% of barcodes were associated with vector sequence outside the T-DNA sequence, indicating integration of unprocessed plasmid into the genome. Approximately 1% of RB-TDNAseq reads mapped equally well to two or more highly similar sequences and thus we could not determine which locus is the true site of insertion. Finally, another 1% of barcodes appeared in distinct RB-TDNAseq reads mapping to two or more sequences, suggesting two or more different mutant strains have received the same barcode, rendering those strains indistinguishable in BarSeq data.

T-DNA can integrate into multiple locations in the same genome, giving rise to confounding phenotypes between different mutations. Rates of multi-locus insertion range widely (5% to 45%) depending on transformation conditions and the targeted cell type (7, 10-13). Multi-locus insertions can be derived from multiple copies of T-DNA from a single transformation event, or from co-transformation of distinct T-DNAs. Since only 1% of barcodes mapped to multiple locations, we inferred the former scenario was rare. To estimate the frequency of multiple insertion events from co-transformation, we isolated single colonies and then sequenced their barcodes using PCR amplification with Sanger sequencing. Of 58 colonies with unambiguous sequence of the common sequence preceding the random barcodes, 41 colonies (71%) had a single, unique sequence in the barcode region, suggesting a single barcode was present, and 17 colonies (29%) had mixed signals in the barcode region suggesting T-DNAs with multiple barcodes were present (example traces in Figure 2 – figure supplement 2B). This estimate may be biased by sequence artifacts and should be taken as an upper bound. Furthermore, co-transformed T-DNAs are often integrated into a single concatemeric repeat (11, 14, 15). Thus, far fewer than 29% of strains may actually harbor T-DNA insertions at multiple loci. Conversely, T-DNA insertions have been shown to cause other local mutations (6% of insertions were associated with deletions of more than 100 bp and 0.7% with local inversions in *A. thaliana*(16)). These combined sources of confounding phenotypes highlight the importance of integrating data from multiple T-DNA insertions in any fitness analysis. As such, our main concern in constructing our mutant pool was to effectively probe the entire genome with multiple inserts per gene.

### Fine-scale biases in T-DNA insertion sites

On a genome level, there was no significant bias in rates of T-DNA insertion, with insertion number proportional to scaffold length (Figure 1 – figure supplement 3A) and no apparent bias in insertion rates with respect to local GC content (Figure 1 – figure supplement 3B). We did observe some bias in T-DNA insertion sites at the kilobase scale, however. T-DNAs were mapped within intergenic regions at a higher rate than expected given the composition of the genome (Figure 1 – figure supplement 3C). For instance, 20% of T-DNA inserts were mapped in promoter regions, even though these regions only constitute 8% of the genome. This bias towards promoter regions is consistent with observations in *Cryptococcus neoformans* (17), in *Magnaporthe oryzae* (12), and with the fact that 41% of T-DNA insertions in S. *cerevisiae* mapped in intergenic regions (18) though only 27% of the S. *cerevisiae* genome is intergenic (19). We also observed further fine-scale variation in the density of mapped insertions, with dozens of T-DNA ‘hotspots’ on each scaffold with a higher local density of T-DNA insertion that cannot be explained by a simulated random integration with the observed biases towards promoters, terminators and five-prime UTRs (Figure 1 – figure supplement 3D). We have not explored the mechanism of these fine scale biases, though microhomology to T-DNA borders and local DNA bendability have been suggested as influencing T-DNA insertion into eukaryotic genomes (12, 20).

### Additional information on calculating fitness scores and T-like test statistics

For each barcoded T-DNA insertion, we calculate the log2 ratio of abundance before and after competitive growth in the experimental condition. F is the average of those ratios (weighted by sequence depth) for all the insertions disrupting a given gene. T is a modified student’s T-statistic, a measure of statistical significance of F that incorporates consistency between individual insertions across biological replicate cultures. We observed a wide range in relative abundance of individual mutant strains (i.e. relative counts for different barcodes in BarSeq data). In a typical fitness experiment, we sequenced each sample to a depth of 20 million reads (as opposed to 900 million reads to map insertion locations by RB-TDNAseq). At this depth, approximately 40,000 mapped barcodes (14%) were too rare to count. Countable barcodes ranged from 1 to 1000 counts per sample with a mode around 10 (Figure 2 – figure supplement 1A). Further, for estimating strain abundance in fitness experiments we considered only insertions in the central 80% of the coding region to avoid confounding data from incorrectly predicted gene boundaries, functional truncated proteins, and altered expression of neighboring genes. Within these constraints, we were able to measure fitness for 6,558 genes (92% of non-essential genes) by tracking abundance of 68,021 insertions in coding regions with a median of 7 insertions per gene (Figure 2 – figure supplement 1B).

### Methionine and Arginine biosynthesis in *R. toruloides*

Our fitness data were consistent with established models of arginine biosynthesis in S. *cerevisiae*, and cysteine and methionine synthesis in *A. nidulans* (Figure 2 – figure supplement 2). Out of 13 genes required to produce methionine from sulfate, one gene (*MET7*) was essential in mutant construction conditions, 11 genes (*MET1, MET2, MET3, MET5, MET6, MET10, MET12, MET13, MET14, MET16, and GDH1*) had significantly different fitness scores between supplemented conditions (YPD, DOC, or methionine) and the non-supplemented condition. One gene (*MET8*) fell just below our statistical cutoffs, with fitness scores suggesting methionine/cysteine auxotrophy, but the magnitude of the T-statistics for supplemented conditions (YPD, DOC, or methionine supplementation) versus non-supplemented conditions never exceeded 2.7. We also noted that though the transulfuration pathway and *MET17* were dispensable, *RTO4_15248* and *RTO4_12031* (orthologs of *A. nidulans cysA* and *cysB*) were required for robust growth, suggesting sulfur uptake occurs primarily through cysteine. Nine of nine genes expected to be required for arginine biosynthesis *(ARG1-8, CPA1*, and *CPA2*) had significant fitness scores suggesting auxotrophy. So did *IDP1* and *GDH1*, suggesting the primary source of glutamate in our conditions was from ammonia and 2-oxoglutarate. The mitochondrial ornithine transporter *ORT1* was also required for arginine prototrophy, but *AGC1* was not, suggesting alternative routes for glutamate transport.

### K-means clusters of fitness scores on fatty acids

Cluster FA1 consists of 21 genes for which mutants had consistent growth defects across all three fatty acids. These genes included three mitochondrial beta-oxidation enzymes; the acyl-CoA dehydrogenase *RTO4_14070* (ortholog of *Homo sapiens ACADSB*), the enoyl-CoA hydratase *RTO4_14805* (ortholog of *H. sapiens ECHS1*), and the hydroxyacyl-CoA dehydrogenase *RTO4_11203* (ortholog of *H. sapiens HADH*). Also included were the electron-transferring-flavoprotein subunit *AIM45* and the electron-transferring-flavoprotein dehydrogenase *CIR2*, likely reflective of electron-transferring-flavoproteins’ known role as an electron receptor for acyl-CoA dehydrogenases (21). The carnitine O-acetyltransferase *CAT2* (involved in fatty-acyl-CoA transfer in the mitochondria (22)) and *PEX11* (involved in peroxisome division and possibly interaction between peroxisomes and mitochondria(23)) were also in cluster FA1. Rounding out cluster FA1 were nine genes with likely roles in gluconeogenesis, glucose homoeostasis and/or growth on non-preferred carbon sources (*FBP1, RTO4_14162* (ortholog of *ICL1*), *MLS1, GLG1, MRK1, SNF1, SNF3, SNF4, RTO4_11412* (similar to *SWI1*); two genes involved in mitochondrial amino acid metabolism (*PUT2 and AGC1*); the peroxidase *RTO4_10811* (ortholog of *CCP1*); and *RTO4_12955*, a LYRM domain-containing protein with likely roles in mitochondrial electron transport (24).

Clusters FA2 through FA7 were comprised of 108 genes for which mutants had stronger fitness defects on one or two fatty acids, primarily genes with stronger defects on methylricinoleic acid and ricinoleic acids (FA2 and FA3, 55 genes) or on ricinoleic acid only (FA4 and FA5, 30 genes). These clusters were comprised of genes with predicted roles in various aspects of cellular homeostasis including amino acid metabolism, glycogen metabolism, phospholipid metabolism, protein glycosylation, the mitochondrial electron transport chain, and 17 genes with no well-characterized homologs. See Supplementary file 2 for a complete list. Clusters FA2 and FA7 also included 10 genes predicted to play direct roles in peroxisomal beta-oxidation, however. Cluster FA2 (stronger defect on methylricinoleic and ricinoleic acid) included *RTO4_10408* (ortholog of *H. sapiens ACAD11*), *RTO4_14567* (similar to *H. sapiens ACAD11*), acyl-CoA oxidase *RTO4_12742* (ortholog of *POX1*), and *RTO4_8673* (similar to *PEX11*). Cluster FA7 (stronger defect on oleic acid) included 3-ketoacyl-CoA thiolase *RTO4_13813* (ortholog of *POT1*), enoyl-CoA hydratase *RTO4_11907* (ortholog of *H. sapiens ECH1*), 3-hydroxyacyl-CoA dehydrogenase/enoyl-CoA hydratase *FOX2*, predicted acyl-CoA dehydrogenase *RTO4_8963*, and peroxisomal signal receptors *PEX7 and RTO4_13505* (similar to *PEX5*).

### BODIPY 493/503 and buoyancy as measures of lipid content in *R. toruloides*

Under carbon-replete growth conditions in which nitrogen, sulfur, or phosphorus are limiting, *R. toruloides* accumulates up to 70% of its dry weight in neutral lipids (25-28). These lipids are stored as triacylglycerides (TAG) in specialized organelles called lipid droplets (reviewed in (29-31))(*R. toruloides* lipid droplets visualized in Figure 4A). BODIPY staining has been used extensively to label lipids and we find that in *R. toruloides* cultures, average cellular BODIPY signal correlates well with total fatty acid methyl ester content as quantified using gas chromatography with flame ionization detection (Figure 4 – figure supplement 1). Because lipid droplets have lower density than most cell components, as cells accumulate large lipid droplets, they become more buoyant (Figure 4 – figure supplement 2).

### NADPH production in *R. toruloides*

*R. toruloides* has two predicted malic enzymes, *RTO4_12761* and *RTO4_13917*, which could theoretically provide NADPH for fatty acid synthesis. Their specificities for NAD+ versus NADP+, are unknown but *RTO4_12761* is more closely related to the NADP-specific malic enzyme from *Mucor circinelloides* (32) and Zhu et al. measured increased protein levels in nitrogen-limited conditions (3). Neither gene had significant enrichment scores in our lipid accumulation assays. We mapped very low insertion density in the major enzymes of the pentose phosphate pathway (the primary source for NADPH in *Y. lipolytica* (33)) in our pool, suggesting it was essential in our library construction conditions. As such, the primary source of NADPH in *R. toruloides* remains unconfirmed. Our data are consistent with recent predictions from a simplified metabolic model for *R. toruloides* that during lipid production from glucose, the pentose phosphate pathway should account for greater metabolic flux and NADPH production than malic enzyme (34).

*YEF1* may also increase the supply of NADPH by phosphorylation of NADH, but presumably this reaction could only play a significant role in fatty acid synthesis if NADP+ is efficiently converted to NAD+ for reduction by NAD(+)-dependent enzymes. NADPH phosphatase activity has been observed for inositol monophosphatases of archaea (35), but these activities have not been well explored in fungal species. Alternatively, *YEF1* may be required for efficient lipid accumulation simply because in its absence the total cytosolic NADP(H) concentration is too low for efficient fatty acid synthesis, regardless of the balance between NADP+ and NADPH.

---

**Figure 1 Supplement 1.**
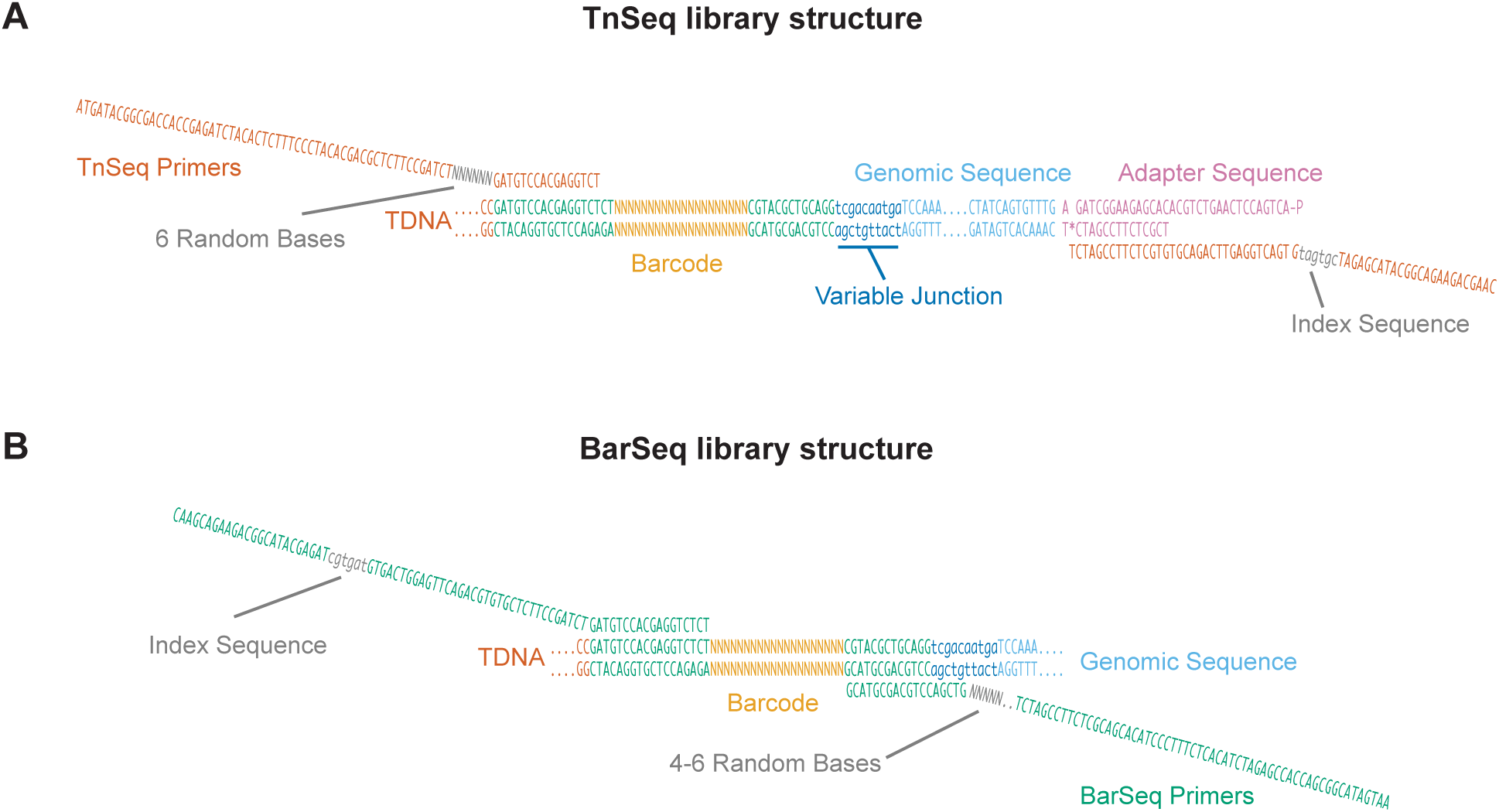
Schematic of TnSeq and BarSeq libraries generated using RB-TDNAseq. (A) In the TnSeq protocol, genomic DNA is sheared into ~300 bp fragments, and Illumina TruSeq adapters are ligated on both ends. T-DNA junctions are then specifically enriched by PCR with a T-DNA-specific and an adapter-specific primer. (B) In the BarSeq protocol, genomic DNA is used as a template for a more robust and quantitative PCR on the barcoded region of the T-DNA insert. Phasing error caused by the identical T-DNA sequences flanking the random barcodes was reduced by adding sequence diversity at the beginning of each read, either by the introduction of a short random 6 bp sequence or a 4-6 bp random sequence for TnSeq and BarSeq, respectively.

**Figure 1 Supplement 2.**
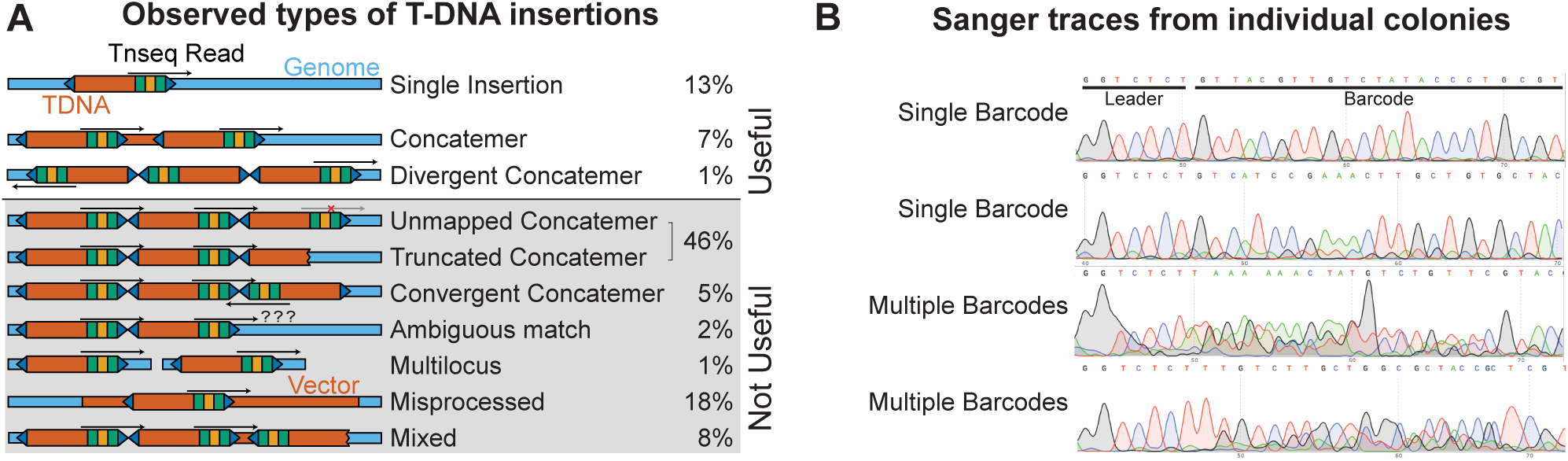
Complexities of T-DNA insertions. (A) Inferred topology of T-DNA insertions from associations of barcodes and adjacent genomic or T-DNA sequence. Only three of the observed insertion types could be mapped using the TnSeq protocol. (B) Sanger sequencing of barcodes from single colonies isolated from the pool. Multiple overlapping peaks in the barcode region suggest multiple T-DNAs are present in a single strain. Note that these T-DNAs may be integrated at the same, or different loci. Inherent noise in barcode amplification and sequencing introduces significant ambiguity in this analysis. The inferred rate of multiple barcode insertion (29%) should be considered a maximum estimate.

**Figure 1 Supplement 3.**
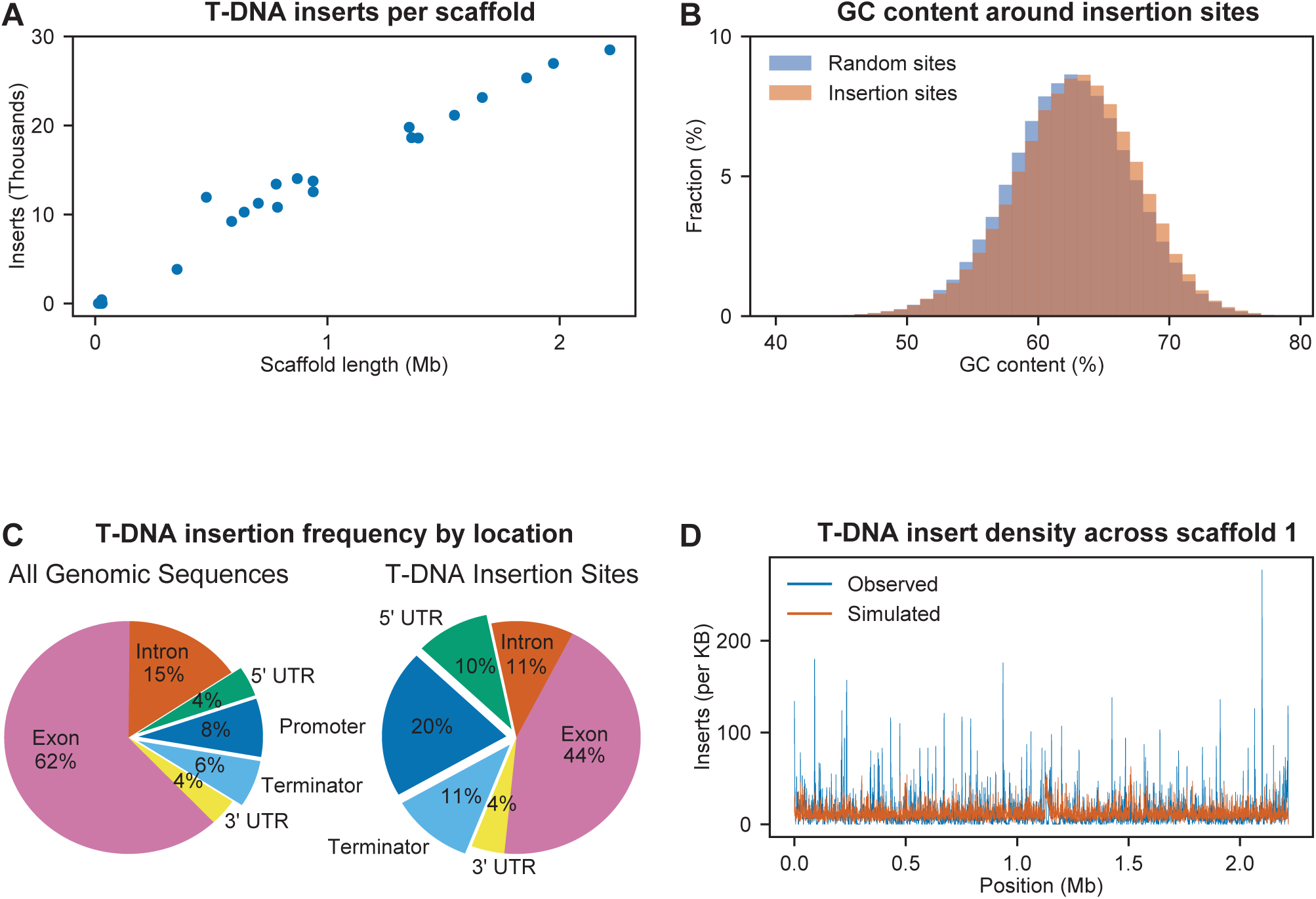
Observed biases in T-DNA insertion locations. (A) Frequency of T-DNA insertion mapping was consistent across all 30 IFO 0880 scaffolds. (B) Histogram of GC content in 100 base pair regions flanking insertion sites and in random 100 base pair regions. (C) Proportion of the *R. toruloides* IFO 0880 genome in promoter regions, terminator regions, untranslated regions transcribed to mRNA, coding exons, and introns versus the proportion of T-DNA insertions mapped to those sequences. (D) Distribution of T-DNA insertion density across the length of scaffold 1. Total inserts were summed across a rolling 1000 base pair window using the observed insertions and a simulated random mutant pool assuming biases for insertion in promoters, terminators and untranslated transcribed regions.

**Figure 2 Supplement 1.**
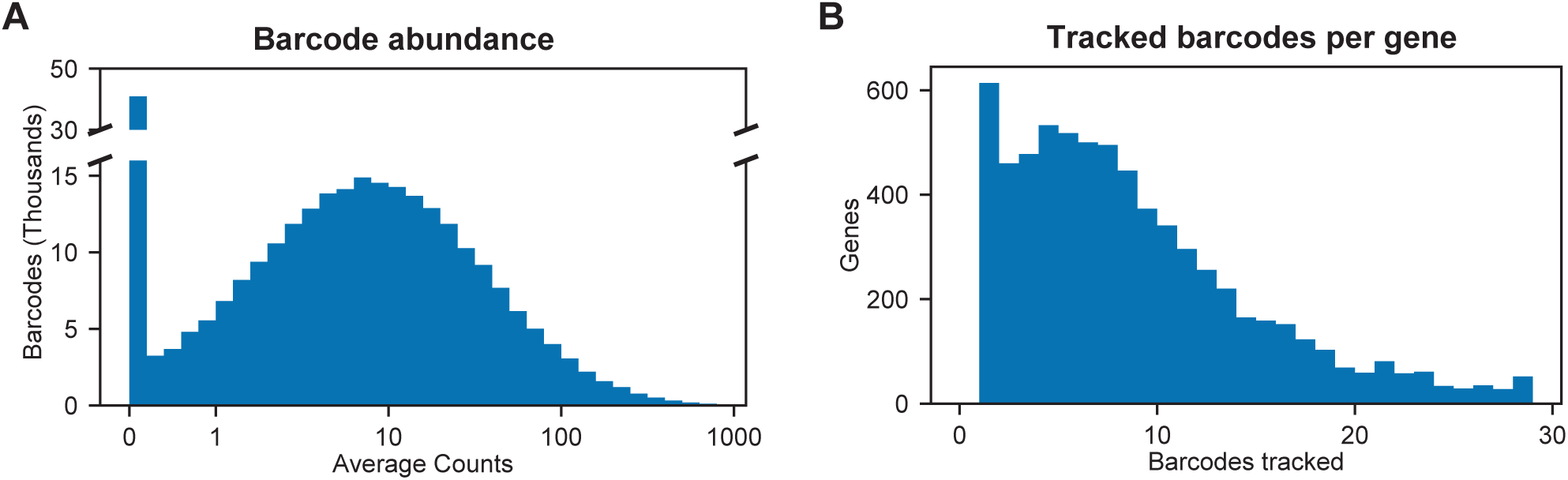
Barcode abundance in BarSeq experiments. (A) Histogram of barcode abundance in a typical BarSeq experiment with 20 million reads per sample. (B) Histogram of tracked barcodes per gene in a typical BarSeq experiment. Median 7 barcodes per gene, 68,021 total barcodes in 6,558 genes. See supplementary file 1 for a full list of insert density by gene and orthologs reported as essential in model fungi.

**Figure 2 Supplement 2.**
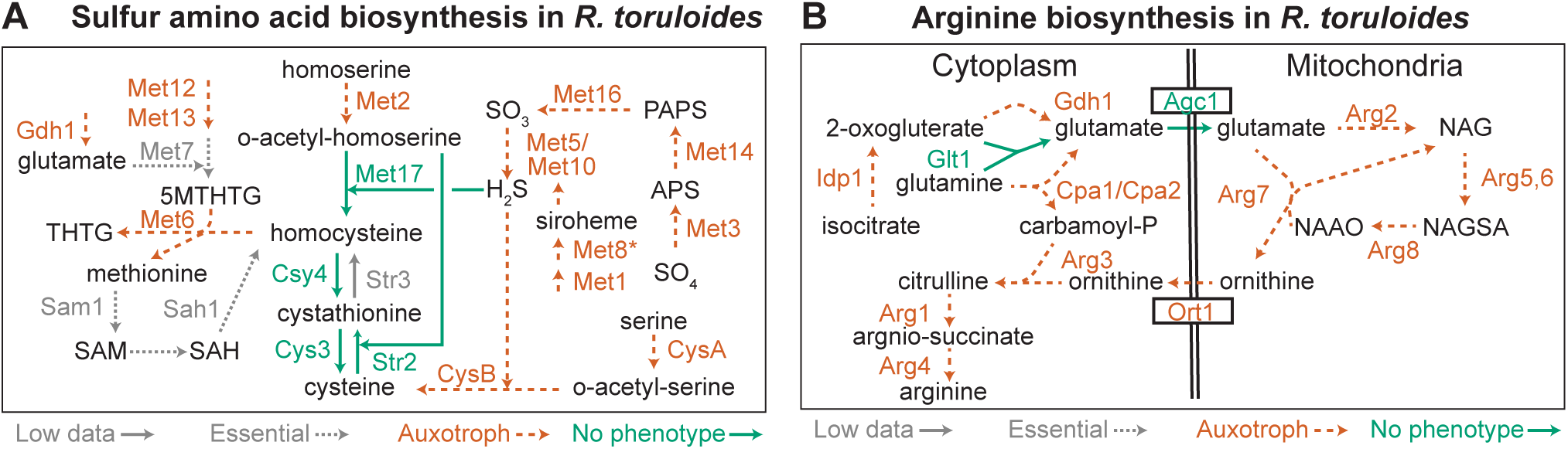
Methionine, cysteine, and arginine biosynthesis pathways in *R. toruloides*. (A) Sulfur amino acid biosynthesis in *R. toruloides* as inferred from enrichment experiments. CysA/CysB are named according to their *A. nidulans* orthologs, all others by orthologs in S. *cerevisiae*. Auxotrophic mutants had F < -1 in non-supplemented media and T < -3 versus the methionine supplementation, drop-out complete or YPD cultures, with the exception of *MET8* which had T < -2. Multiple insertions were mapped in *STR3*, suggesting non-essentiality, but strain abundance was too low to reliably estimate fitness in BarSeq experiments. 5MTHTG: 5-methyltetrahydropteroyltri-L-glutamate, THTG: tetrahydropteroyltri-L-glutamate, SAM: S-adenosyl-L-methionine, SAH: S-adenosyl-homocysteine, APS: adenylyl-sulfate, PAPS: 3'-phosphoadenylyl-sulfate. (B) Arginine biosynthesis in *R. toruloides* as inferred from enrichment experiments. Gene names are based on orthologs in S. *cerevisiae*. NAG: N-acetylglutamate, NAGSA: N-acetylglutamate semialdehyde, NAAO: N-alpha-acetylornithine.

**Figure 3 Supplement 1.**
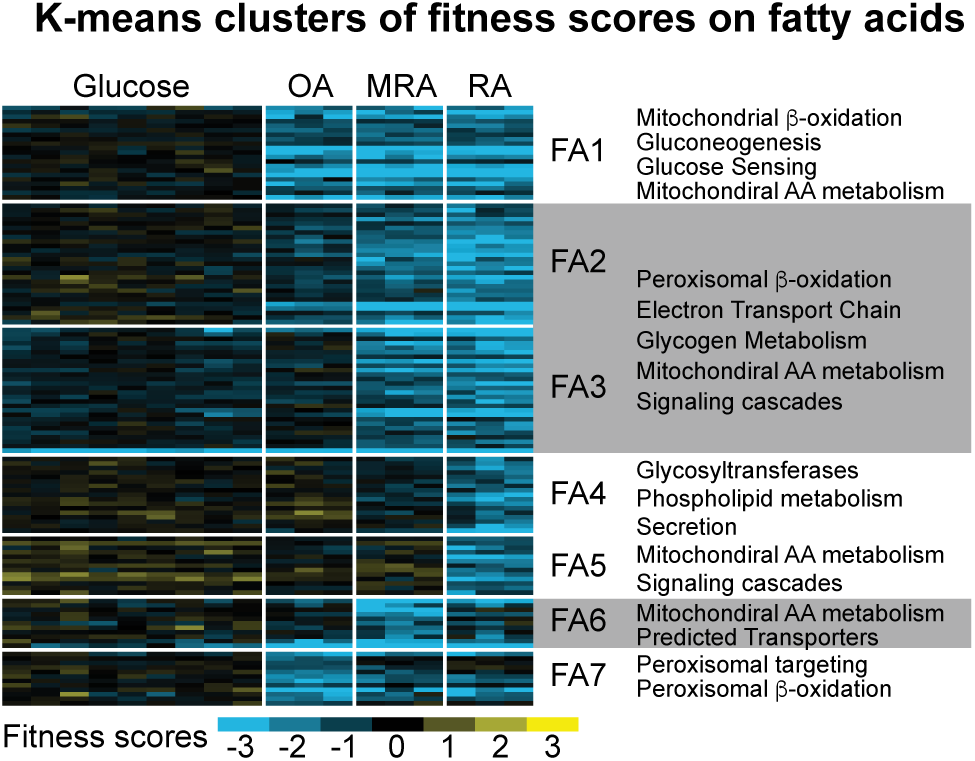
K-means clusters of fitness scores for 129 genes for which mutants have specific fitness defects on fatty acids. Fitness scores for individual biological replicates were clustered in this analysis (6 replicates on glucose, 3 for each fatty acid). OA: oleic acid, RA: ricinoleic acid, MRA: methyl ricinoleic acid. Seven clusters were identified based on carbon utilization patterns; FA1 - fitness defects on all fatty acids, FA2 & FA3 - fitness defects on MRA and RA, FA4 & FA5 – fitness defects on RA only, FA6 – fitness defects on MRA only, and FA7 – fitness defects on OA only. Major categories of predicted gene functions are summarized for the clusters. See supplementary files 2 and 3 for full fitness data and gene ontology enrichments.

**Figure 3 Supplement 2.**
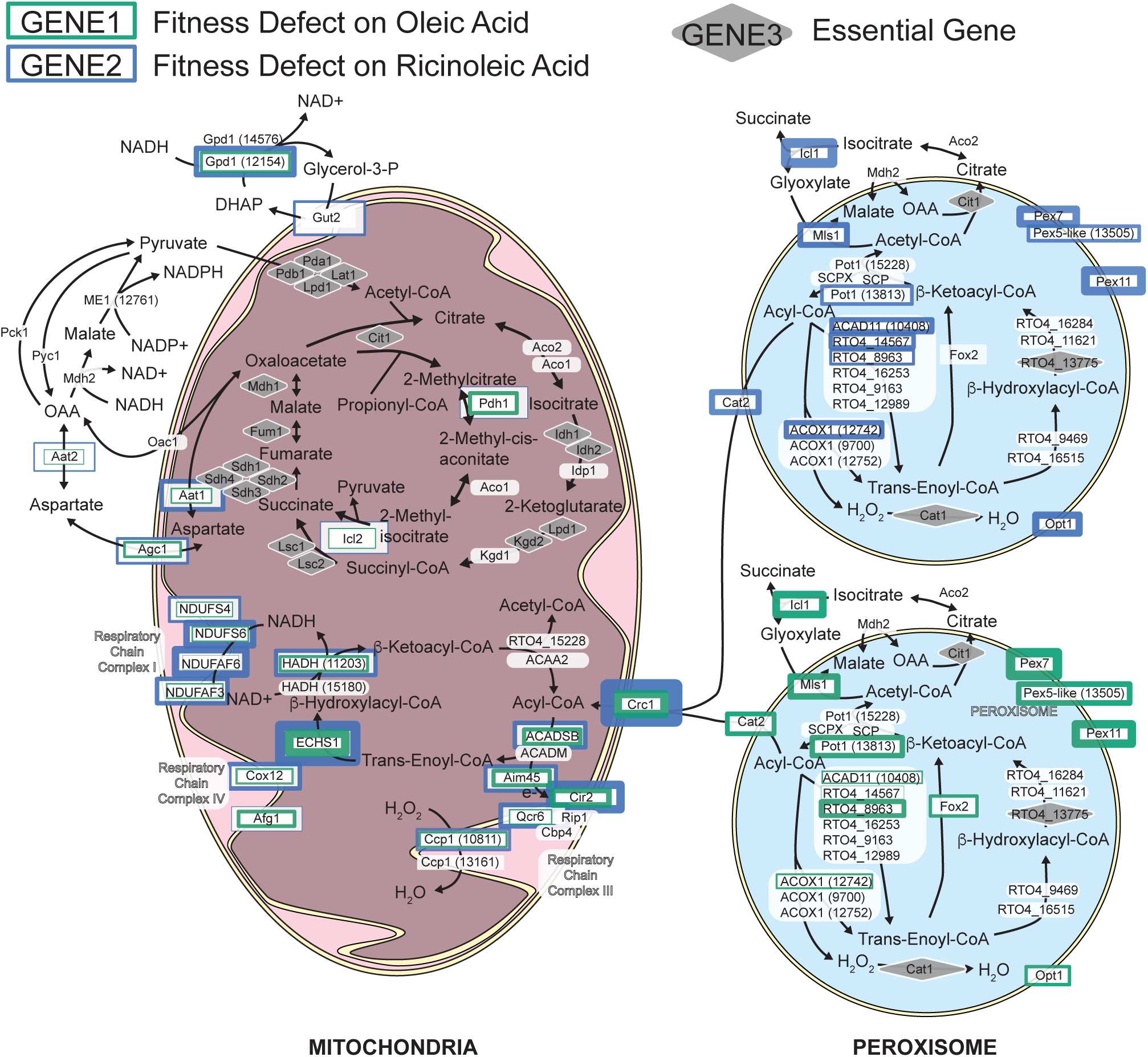
Model for beta-oxidation of fatty acids in *R. toruloides*. Fitness scores for genes with predicted roles in mitochondrial and peroxisomal beta-oxidation are represented by the width of green or blue borders around each protein, with wider borders corresponding to lower fitness scores. Green and blue borders represent fitness on oleic and ricinoleic acid respectively. Fitness scores on fatty acids were consistently most severe for a few mitochondrial beta-oxidation genes, and an ortholog to the mammalian short-chain and branched short-chain acyl-CoA dehydrogenase *ACADSB* was the most important gene mediating that enzymatic step in the mitochondria. Fitness scores were more variable between different fatty acids for peroxisomal enzymes, for which more paralogs are present.

**Figure 3 Supplement 3.**
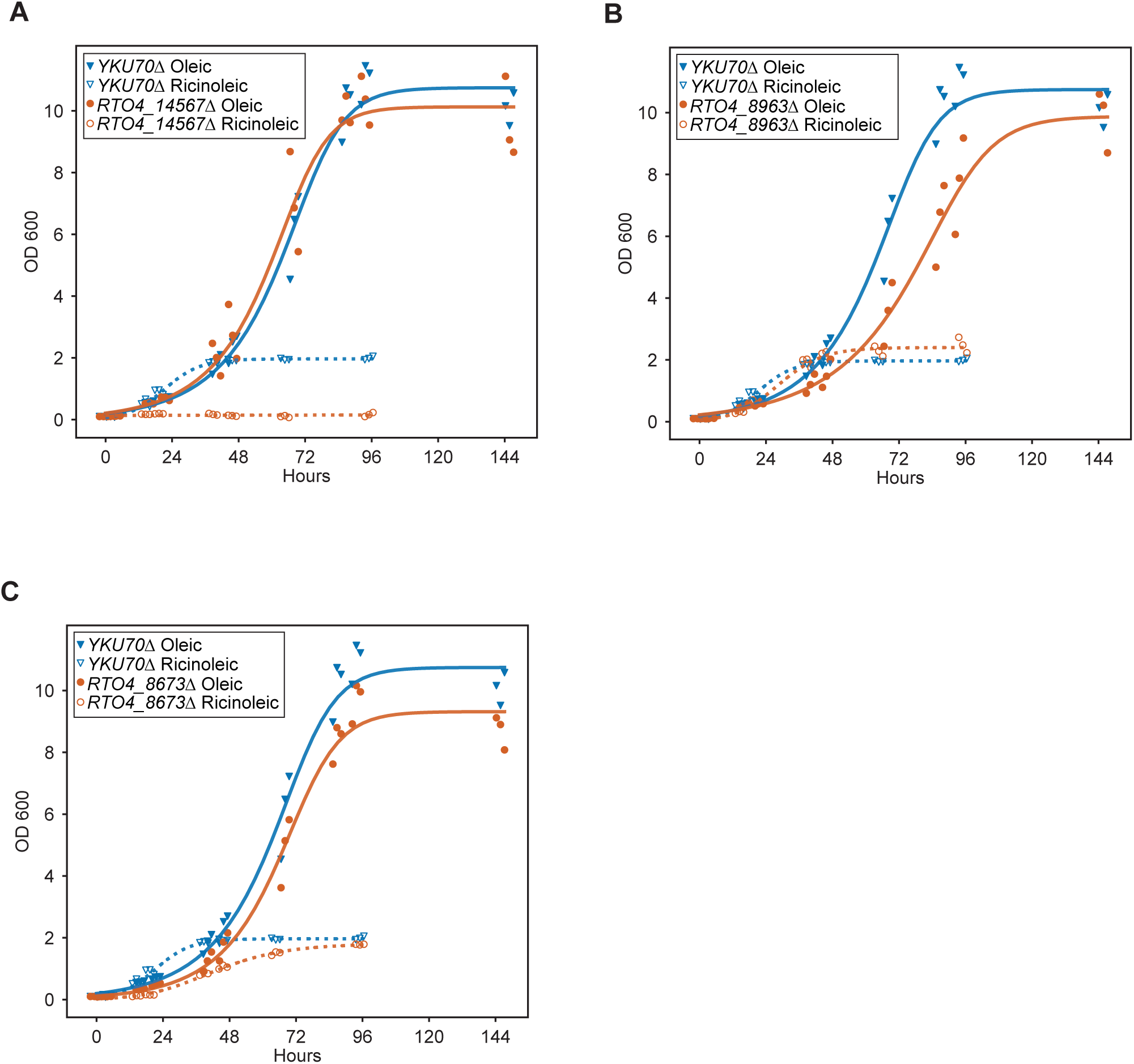
Extended growth curves for deletion mutants on fatty acids. Growth curves for deletion mutants of (A) *RTO4_14567* (similar to *H. sapiens ACAD11*), (B) *RTO4_8963* (similar to *H. sapiens ACAD11*), and (C) *RTO4_8673* (similar to *PEX11*) on 1% oleic acid and 1% ricinoleic acid as the sole carbon source.

**Figure 4 Supplement 1.**
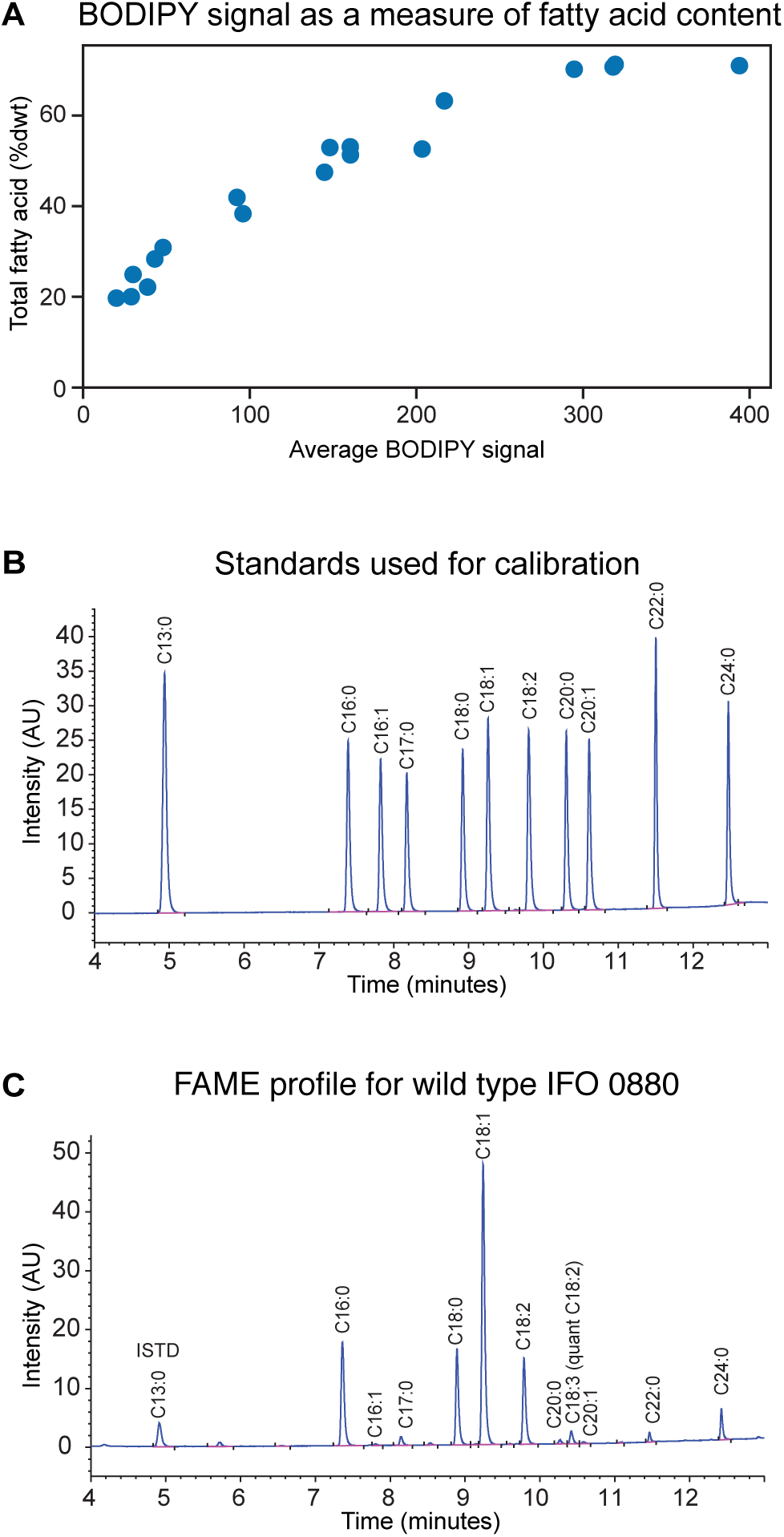
Measuring lipid accumulation under nitrogen limitation. (A) Total fatty acid methyl ester (FAME) content in *R. toruloides* cultures, quantified using gas chromatography and flame ion detection (GC-FID), correlates with average cellular BODIPY signal determined by flow cytometry. (B) Standards used for quantification of FAME content. Peak area/concentration ratios for ten commercially available fatty acid standards were used to quantify FAME peaks from experimental samples. (C) Example FAME profile for IFO 0880. Peak area/concentration ratios for C18:2 were used to quantify C18:3.

**Figure 4 Supplement 2.**
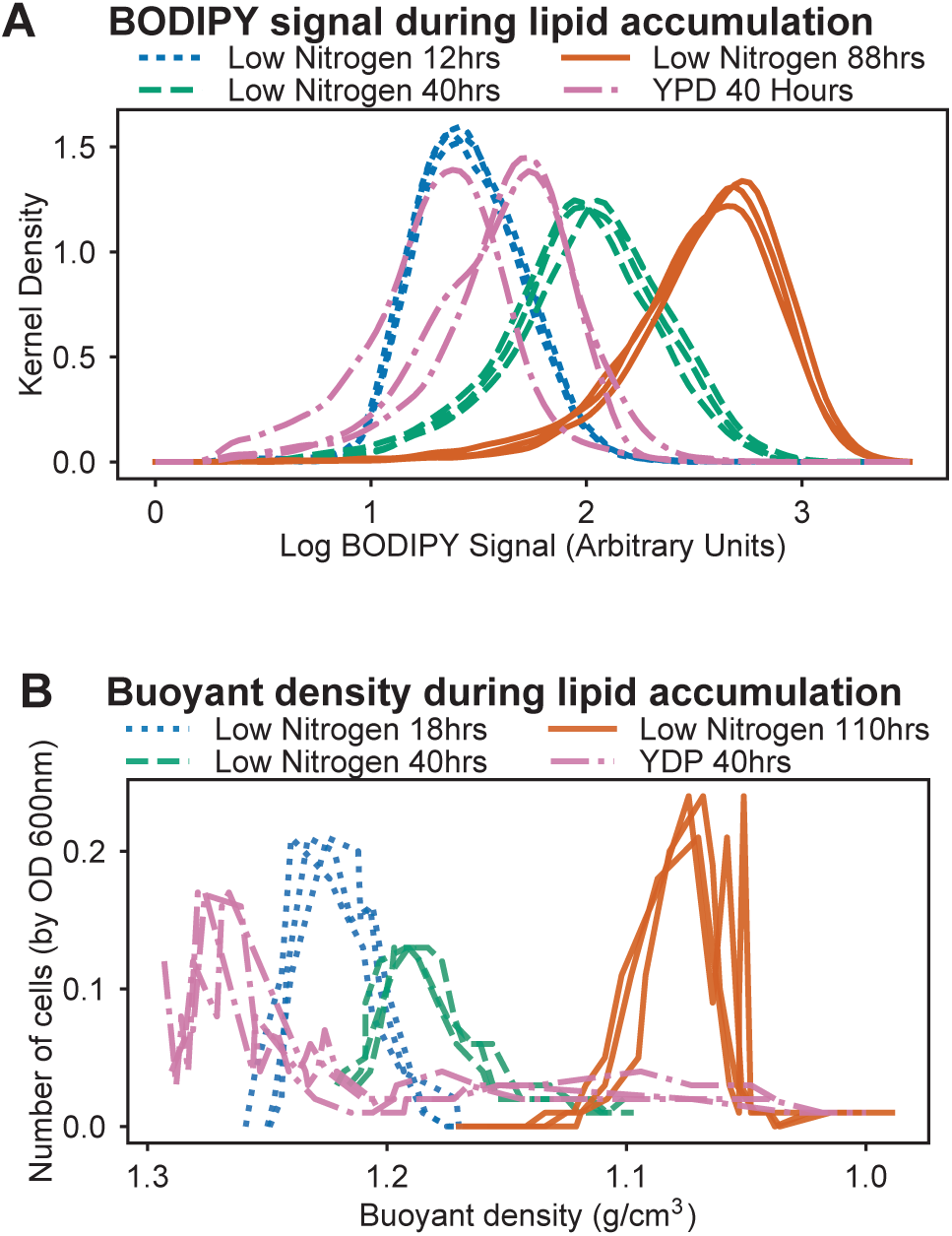
Lipid accumulation and buoyancy changes under nitrogen limitation. (A) Time course of lipid accumulation (measured by BODIPY intensity) in nitrogen limited media (C/N 120; 12, 40, and 88 hours). Rich media control shown for comparison (YPD at 40 hours). Kernel Density plots for three biological replicates are shown for each growth condition. (B) Time course of buoyant density on sucrose gradients in nitrogen limited media (C/N 120; 12, 40, and 88 hours). Rich media control shown for comparison (YPD at 40 hours). Relative cell numbers were measured by OD 600 nm. Density was measured directly by weight of a 100 μl sample.

**Figure 4 Supplement 3.**
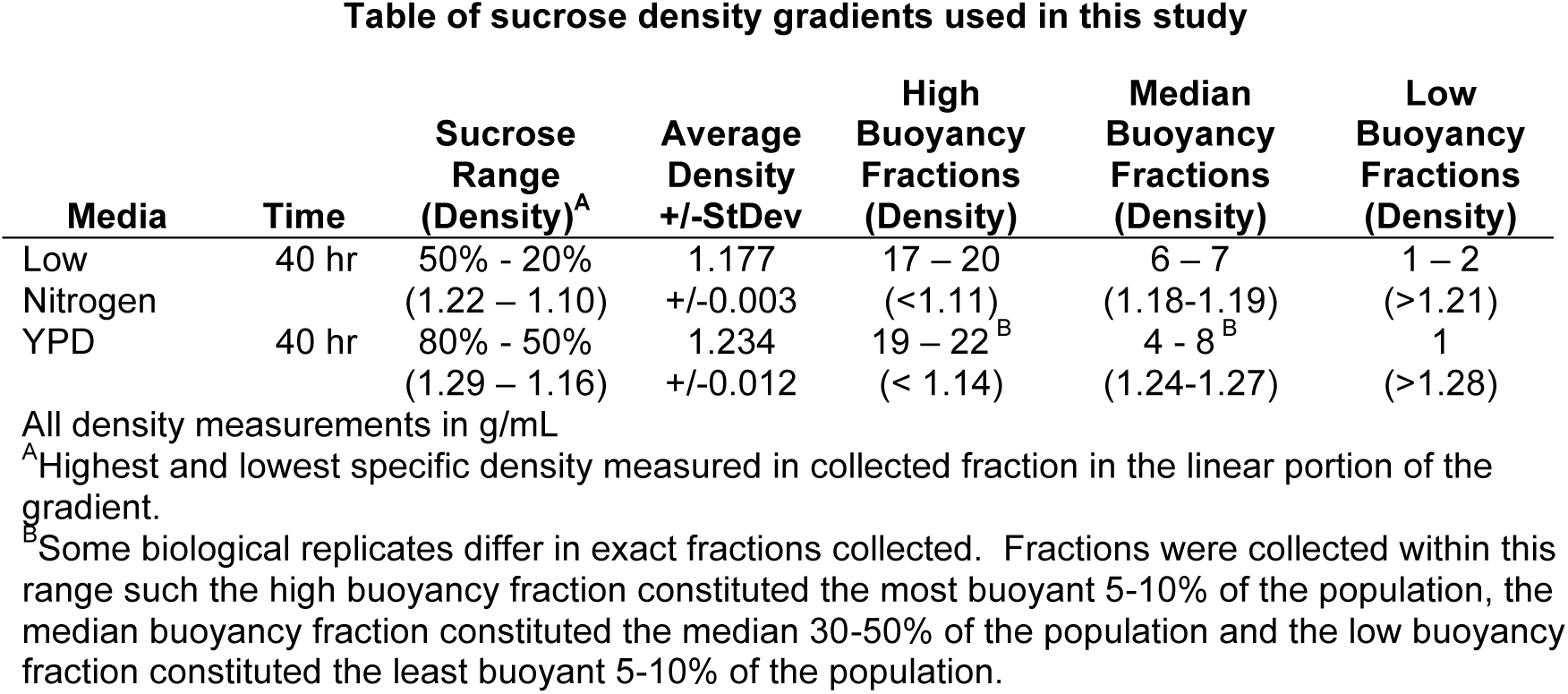
Table of sucrose gradients used in this study.

**Figure 5. Supplement 1.**
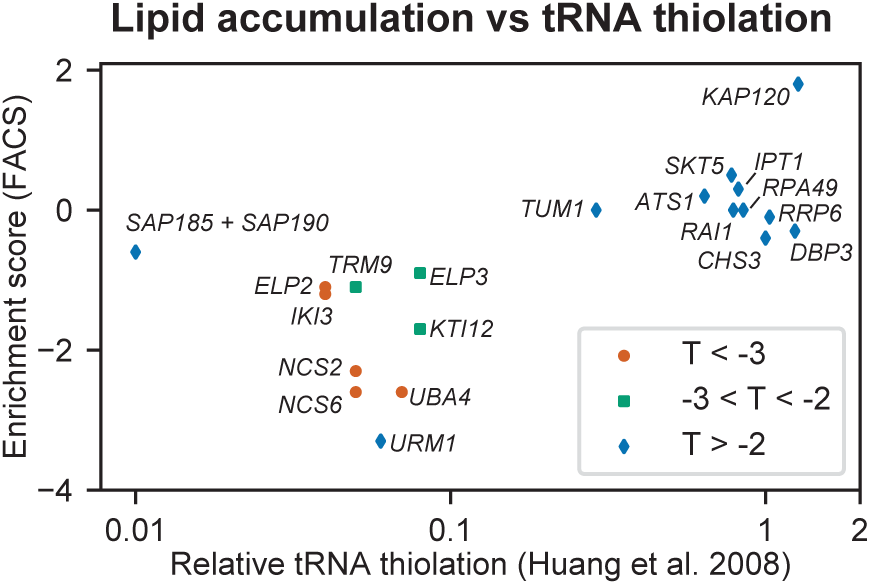
tRNA thiolation in *S. cerevisiae* versus lipid accumulation in *R. toruloides*. Relative levels of tRNA thiolation for *S. cerevisiae* mutants as reported by Huang et al(22) versus enrichment scores for orthologous *R. toruloides* genes in the FACS separation experiment after lipid accumulation. Low lipid content (i.e. negative enrichment scores) for *R. toruloides* mutants corresponds to lower levels of tRNA thiolation in *S. cerevisiae* mutants.

**Figure 6. Supplement 1.**
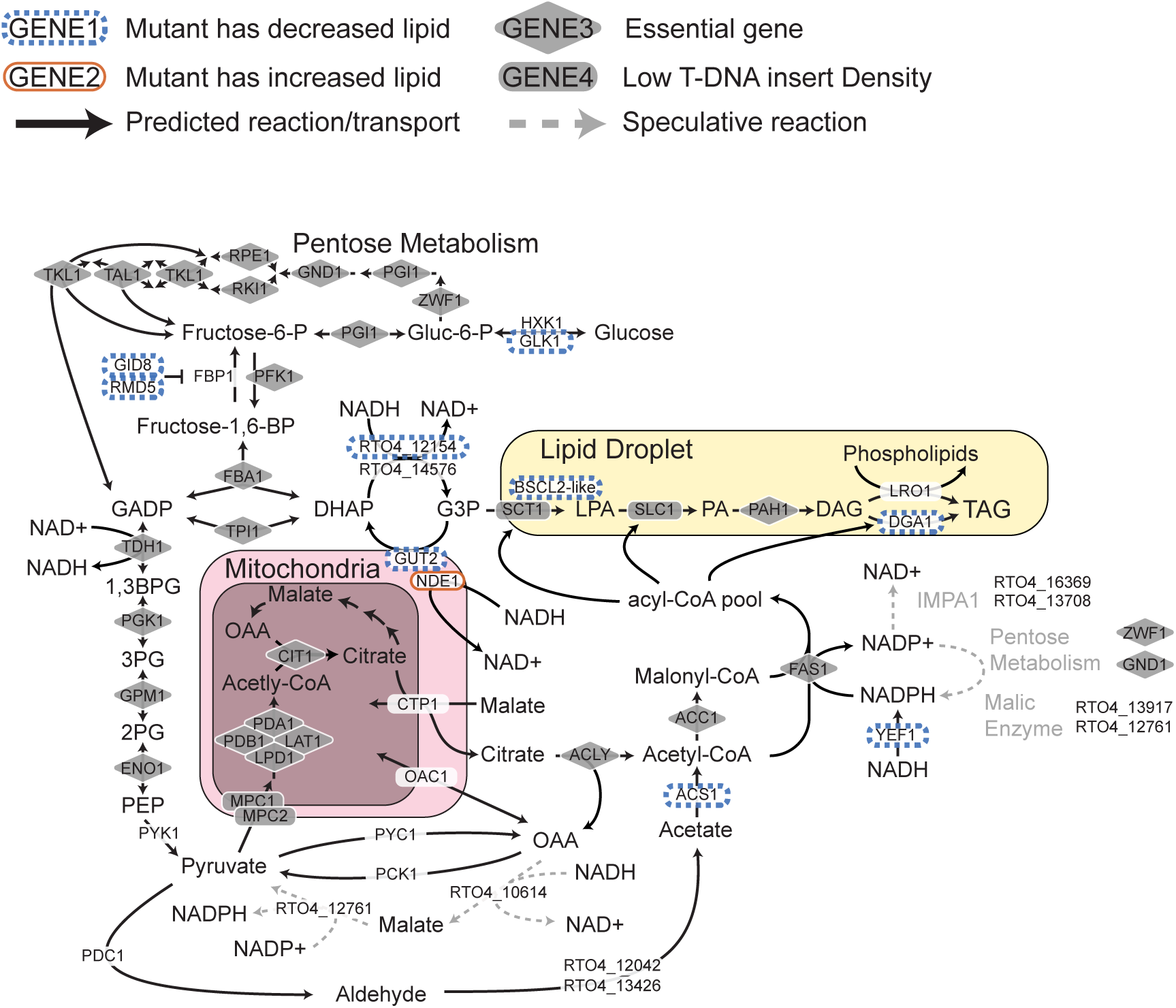
Genes directly effecting TAG biosynthesis in *R. toruloides*. Model pathway illustrating genes involved in glycolysis, triacylglyceride (TAG) synthesis, and cytosolic NAD+/NADH balance during TAG synthesis. Genes for which mutants had altered lipid accumulation (enrichment scores in clusters LA1, LA6, LA7, or LA8) are highlighted in orange or blue. Genes with low rates of T-DNA insertion (essential genes and genes for which mutants have a strong growth defect) are highlighted in gray. The primary source of NADPH in *R. toruloides* remains unclear (see supplementary text for detail). Speculative pathways mediating NADPH production are indicated with dashed gray arrows. DaG: diacylglycerol, PA: phosphatidic acid, LPA: lysophosphatidic acid, G3P: glycerol-3-phosphate, DHAP: dihydroxy-acetone-phosphate, GADP: glycerate 3-phosphate, 1,3BPG: 1,3-bisphosphoglycerate, 3PG: 3-phosphoglycerate, 2PG: 2-phosphoglycerate, PEP: phosphoenolpyruvate, OAA: oxaloacetate

**Figure 7 Supplement 1.**
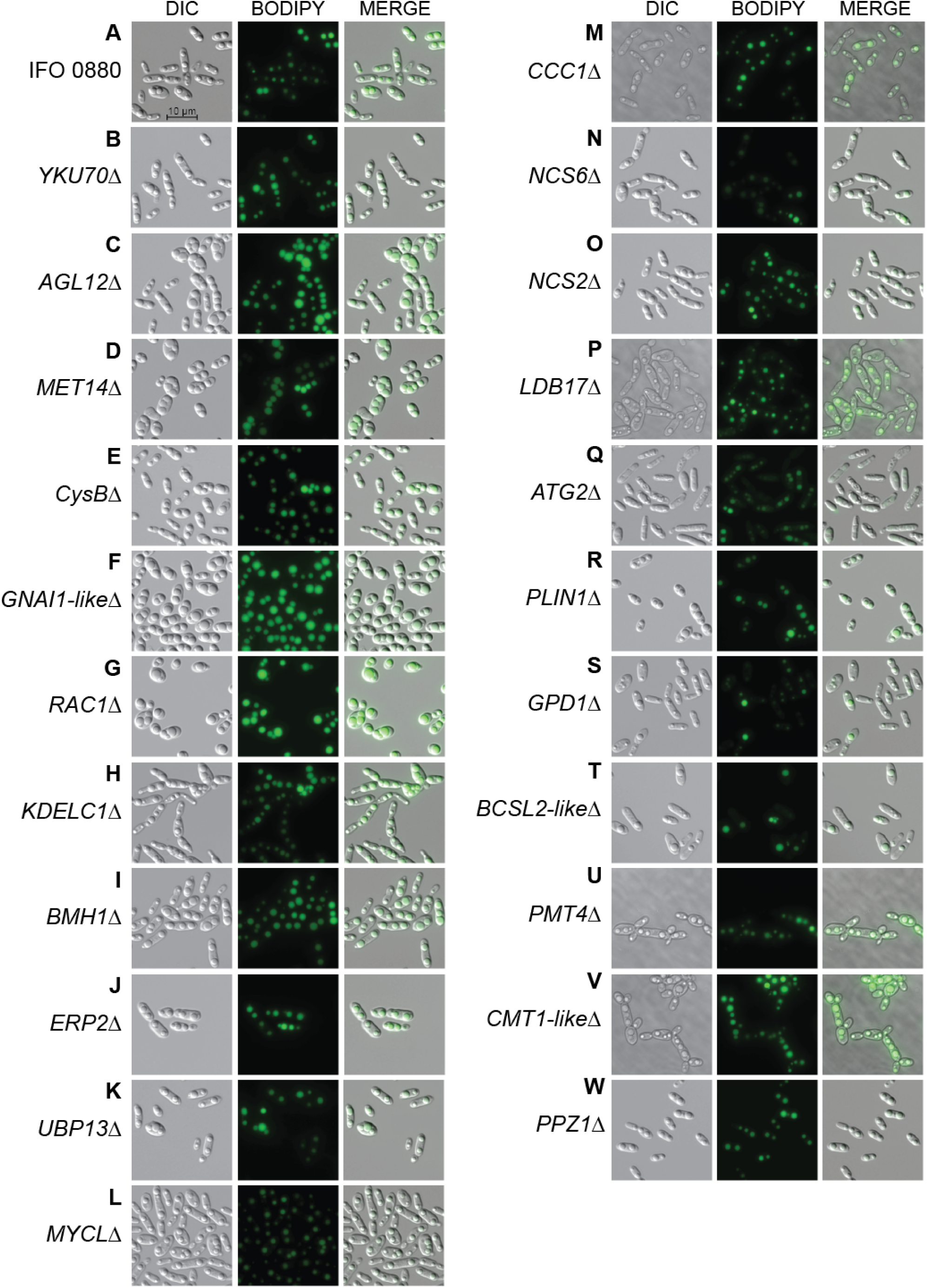
Additional light and fluorescence microscopy images. DIC microscopy on 21 deletion mutants for lipid accumulation genes. All deletion mutants (C-W) were constructed in a *YKU70*Δ background to enable homologous recombination at the targeted locus. Cells were grown 40 hours in low nitrogen lipid accumulation media. DIC, BODIPY 493/503 fluorescence, and composite images are shown for 23 strains. (A) *R. toruloides* IFO 0880 (WT). (B) *RTO4J1920*Δ ortholog of *YKU70*. (C) *RTO4_11272*Δ ortholog of *ALG12*. (D) *RTO4J8709*Δ ortholog of *MET14*. (E) *RTO4_12031*Δ ortholog of *A. nidulans CysB*. (F) *RTO4_16215*Δ similar to *H. sapiens GNA1*. (G) *RTO4_14088*Δ ortholog of *H. sapiens RAC1*. (H) *RTO4_10371*Δ similar to *H. sapiens KDELC1*. (I) *RTO4_16644*Δ ortholog of *BMH1*. (J) *RTO4J 6731*Δ ortholog of *ERP2*. (K) *RTO4_9026*Δ ortholog of *UBP13*. (L) *RTO4_15890*Δ similar to *H. sapiens MYCL*. (M) *RTO4_8506*Δ ortholog of *CCC1*. (N) *RTO4J2817*Δ ortholog of *NCS6*. (O) *RTO4_10764*Δ ortholog of *NCS2*. (P) *RTO4_9970*Δ ortholog of *LDB17*. (Q) *RTO4_13598*Δ ortholog of *ATG2*. (R) *RTO4_16381*Δ similar to *H. sapiens PLIN1*. (S) *RtO4_12154*Δ ortholog of *GPD1*. (T) *RTO4_11043*Δ similar to *H. sapiens BSCL2*. (U) *RTO4J2121*Δ ortholog of *PMT4*. (V) *RTO4_10302*Δ similar to *C. neoformans CMT1*. (W) *RTO4_11380*Δ ortholog of *PPZ1*.

